# Exosomes are specialized vehicles to induce fibronectin assembly

**DOI:** 10.1101/2025.06.23.661091

**Authors:** Bong Hwan Sung, Merlyn Emmanuel, Metti K. Gari, Jorge F. Guerrero, Maria Virumbrales-Muñoz, David Inman, Evan Krystofiak, Alan C. Rapraeger, Suzanne M. Ponik, Alissa M. Weaver

## Abstract

Fibronectin is a key stromal matrix molecule whose assembly into fibrils is thought to require cells. In contrast, we find that small exosome-type extracellular vesicles (EVs) are critical initiators of fibronectin assembly. Fibroblasts engineered to be deficient in exosome secretion showed greatly reduced assembly of fibronectin and other stromal matrix molecules in 2D, 3D, and *in vivo* environments, and led to reduced tumor growth and lung metastasis by breast cancer cells. Furthermore, transgenic mice with defects in exosome secretion had greatly reduced lung fibrosis after treatment with bleomycin. In a direct test of exosome function, we find that the addition of purified small EVs to purified soluble fibronectin in a cell-free system is sufficient to induce fibronectin assembly. The EV-induced fibronectin assembly requires the presence of fibronectin-binding integrins and Syndecan-1 in the EVs. We propose a new model in which secreted exosomes directly drive stromal matrix assembly and tissue fibrosis.

## Introduction

Extracellular matrix (ECM) holds cells together in multicellular organisms, provides mechanical strength, and forms tracks that facilitate cellular migration. Stromal matrix is largely produced by tissue fibroblasts, is necessary for tissue repair, scar formation, and cellular migration, and is dysregulated in fibrotic diseases, including cardiac, pulmonary, and kidney disorders, and cancer. As the initiating template molecule for stromal matrix assembly, how fibronectin is assembled has been of great interest. The current paradigm is that soluble fibronectin is secreted from cells and assembled at the cell surface. Interaction of soluble fibronectin with cell surface integrins is thought to lead to integrin clustering, increasing the local fibronectin concentration to allow interaction of fibronectin monomers with each other and assembly into multimeric fibrils^1^. Contraction of actin filaments associated with integrins, as well as the interaction of fibronectin and integrins with syndecan proteoglycans, may facilitate this process by inducing fibronectin unfolding. This model explains the proposed requirement for cells in matrix assembly since integrins that bind fibronectin are transmembrane adhesion molecules. Thus far, no acellular factor has been described that can initiate this complex process.

Extracellular vesicles (EVs) are small lipid-enclosed particles that are actively released from cells and carry bioactive protein, lipid, and nucleic acid cargoes ^2^. Cells release several types of EVs, including ectosomes shed from the cell surface and exosomes secreted from late endosomal multivesicular body (MVB) compartments. Ectosomes can be either large (>150 nm) or small (<150 nm) in size, whereas exosomes are small EVs formed as intralumenal vesicles within MVBs^3^. Over the past decade, it has become clear that EVs are released from diverse cell types and organisms and play a fundamental role in autocrine and paracrine cell communication.

Integrins and ECM molecules are frequently identified in EV samples^4–8^ and have been implicated in driving cell migration and cancer metastasis. We found that fibronectin association with exosomes depends on the presence of α5 integrin and that exosome-associated fibronectin drives nascent adhesion formation and cell migration by cancer cells^4,7^. Those data suggested that not only is fibronectin a true EV cargo but that it is present in an assembled, adhesive form. Based on these data, we hypothesized that exosomes can directly induce fibronectin assembly. We tested this hypothesis using multiple model systems, including genetic inhibition of exosome secretion in stromal cells and assessment of cell-associated matrix in 2D, 3D, and *in vivo*.

Notably, inhibition of exosome secretion by knockdown or knockout of Rab27a and/or Rab27b leads to decreased fibronectin assembly in all systems, including in breast mammary tumors and a bleomycin-induced pulmonary fibrosis model. Loss of exosome secretion in fibroblasts also diminished tumor growth and spontaneous lung metastasis of co-mingled breast cancer cells. Finally, we developed and used a cell-free assay to directly demonstrate the sufficiency of EVs in inducing fibronectin assembly and to dissect the role of key fibronectin-binding EV cargoes.

## RESULTS

### Exosome secretion is critical for the assembly of cell-derived stromal matrices

EV-mediated assembly of fibronectin should result in the embedding of EVs within the matrix. Indeed, immunostaining of cell-derived stromal matrix prepared from hTERT-immortalized human mammary fibroblasts (hMFs)^9^ revealed punctate staining for the exosome marker CD63^10^ along fibronectin fibrils (Fig. 1a). To test whether exosome secretion is important for fibronectin assembly, we knocked down (KD) Rab27a and Synaptotagmin VII (Syt7) (Extended Data Fig. 1a), which respectively regulate the docking and fusion of MVBs with the plasma membrane^11–13^, in hMFs. Indeed, Rab27a and Syt7 KD greatly reduced the secretion of small EVs, which represents an exosome-enriched EV population, but not of large EVs (Extended Data Fig. 1b). To test whether exosome secretion affects stromal matrix assembly, cell-derived stromal matrix was prepared from control, Rab27a, and Syt7 KD fibroblasts.

**Figure 1.**
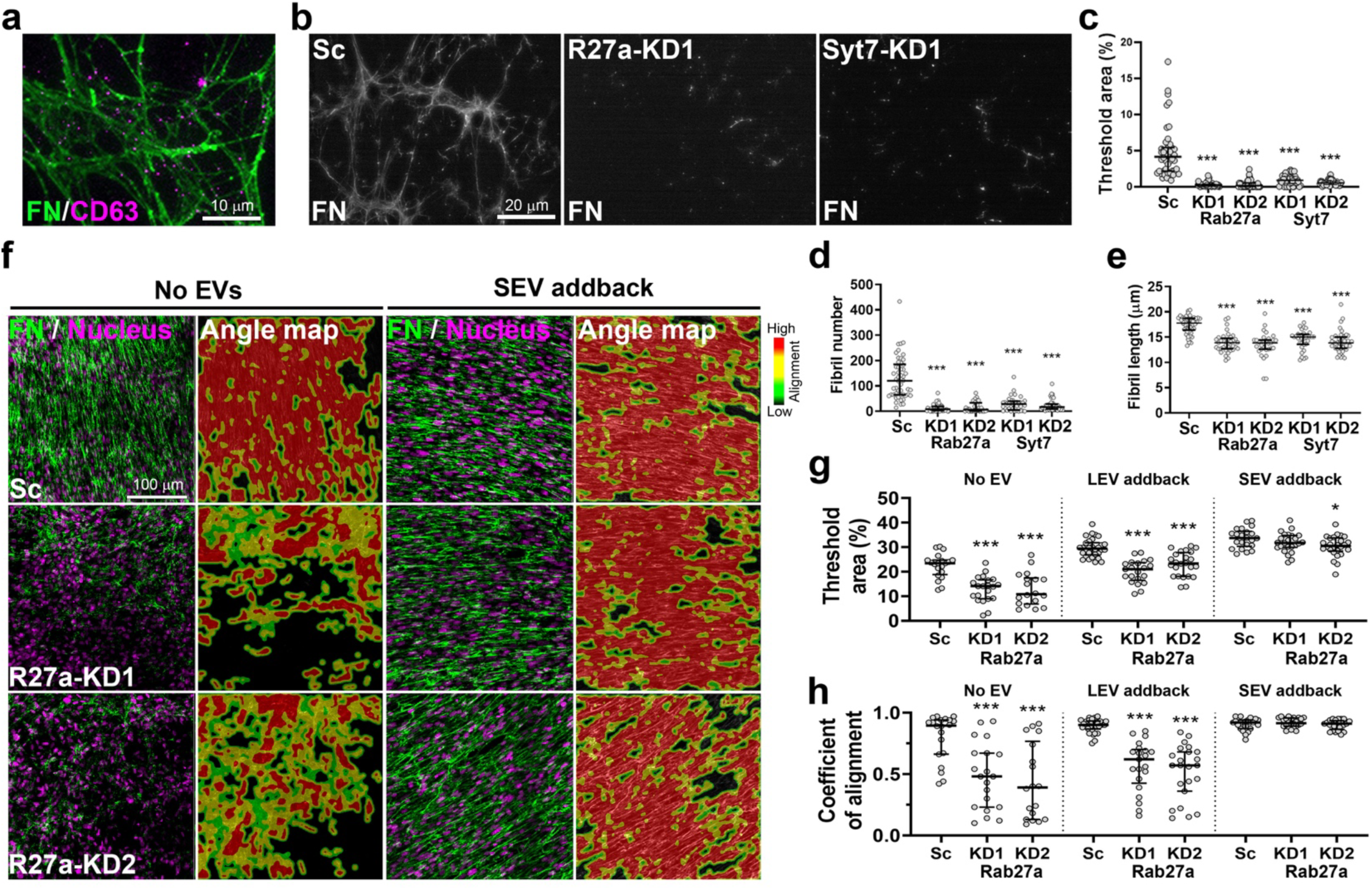
Exosome secretion is critical for stromal matrix assembly and alignment. a, Representative wide field image of cell-derived stromal matrix immunostained for fibronectin (FN) and the exosomal marker CD63. From n=7 images from 1 experiment **b,** Representative images of fibronectin immunostaining on cell-derived stromal matrix prepared from scrambled control (Sc), or Rab27a (R27a)- or synaptotagmin 7 (Syt7)-knockdown (KD) hMFs. **c – e,** Integrated intensity (**c**), fibril number (**d**), and fibril length (**e**) of fibronectin fibrils from **b** (median with interquartile range). n ≥ 32 images from 3 independent experiments. **f,** Representative confocal images by maximum intensity projection of FN immunostaining from Sc or Rab27a-KD hMFs cultured in collagen gels without EVs, or with purified small EVs (SEV) for 7 days. **g** and **h,** Threshold area (**g**) and coefficient of alignment (**h**) of fibronectin fibrils from **f** (median with interquartile range). n ≥ 18 images from 4 independent experiments. *, *p* < 0.05; ***, *p* < 0.001 compared to Sc in the same EV condition by Mann-Whitney *U* test.

Western blot and immunofluorescence analyses of the resulting cell-derived matrix revealed that Rab27a and Syt7 KD cells deposited less fibronectin and tenascin C than control cells (Fig. 1b and c-e, Extended Data Fig. 1c and d). Image analysis revealed that both fibril number and length were reduced in samples produced from exosome secretion-inhibited cells (Fig. 1d and e). In addition, total internal reflection fluorescence imaging of fixed, permeabilized control and KD cells immunostained for fibronectin revealed greatly reduced levels of fibronectin at the cell-substrate interface of KD cells (Extended Data Fig. 1e and f). Western blot analysis of conditioned media collected from control and KD cells over 48 h revealed that there is no significant difference in the amount of secreted soluble fibronectin (Extended Data Fig. 1g), indicating that the difference in fibronectin incorporation into cell derived matrix is due to alterations in fibronectin assembly rather than in synthesis or secretion. Analysis of autocrine-produced stromal matrix from two other stromal cell types, cancer-associated fibroblasts and cardiac stromal cells, validated our finding that inhibition of exosome secretion reduces the assembly of cell-derived stromal matrix (Extended Data Fig. 1h-o).

### Exosome secretion promotes stromal matrix assembly and alignment in 3D cultures

Fibronectin assembly is a precursor to the organization of stromal matrix into aligned fibers, a process associated with fibrosis and cancer metastasis^14–16^. To further test the role of exosomes in this process, control and Rab27a-KD hMFs were embedded in 3D type I collagen gels and cultured for 7 days before fixing and immunostaining for fibronectin and tenascin C, which are stromal matrix molecules associated with the premetastatic niche and cardiac fibrosis^17–19^. Analysis of maximum intensity projection fluorescent confocal images revealed that the levels of fibronectin and tenascin C were significantly reduced in the KD cultures compared to controls (Fig. 1f and g, Extended Data Fig. 2a, e, and f). Analysis of fibril number, length, and alignment further revealed that fibronectin and tenascin C fibrils were less numerous, shorter, and less aligned in the Rab27a-KD cell cultures compared to controls (Fig. 1h, Extended Data Fig. 2b–d, and g-j).

For comparison, we transiently knocked down fibronectin in control hMFs using siRNA (Extended Data Fig. 3a) and conducted the 3D collagen gel assay. As expected, there was a large decrease in the levels of fibronectin in gels containing fibronectin-KD hMFs (Extended Data Fig. 3b-g), which was greater than the decrease observed in Rab27a-KD cultures (69.7% vs. 46.9% reduction, compare Extended Data Fig. 3c to Fig. 1g). Conversely, there was a smaller alteration in fiber alignment in the fibronectin-KD cell gels compared to Rab27a-KD cell gels (20.9% vs. 51.4% reduction, compared Extended Data Fig. 3g to Fig. 1h). Consistent with the known role of fibronectin in templating stromal matrix assembly, tenascin C deposition and alignment was reduced in fibronectin-KD cells (Extended Data Fig. 3h-m), although to a lesser extent than that of Rab27a-KD cell gels (16.8% vs. 69.8% reduction of deposition and 33.8% vs. 50.7% reduction of alignment, comparing Extended Data Fig 3i and m to Extended Data Fig 2f and j, respectively). These data suggest that exosomes are a strong regulator of stromal matrix organization, even beyond fibronectin assembly.

As KD of Rab27a could theoretically also affect cellular functions besides exosome secretion, we tested whether we could rescue KD cell defects in cellular matrix assembly by adding back purified small EVs (SEVs), which represent an exosome-enriched EV preparation. As a negative control, we also tested add-back of large EVs (LEVs), which represent an ectosome-enriched population derived from the plasma membrane and are not affected by Rab27a-KD (Extended Data Fig. 1b). LEVs were isolated by centrifuging hMF conditioned medium for 30 min at 10,000 x g. SEVs were then isolated from the supernatant using a cushioned-density gradient ultracentrifugation method that minimizes EV aggregation and improves separation from protein aggregates^20^. SEVs were characterized by immunoblotting, nanoparticle tracking analysis (NTA), and transmission electron microscopy (TEM) (Extended Data Fig. 4a-c), consistent with current standards ^3^. As expected, purified SEVs efficiently rescued Rab27a-KD cell defects in fibronectin fibril number, length, and organization (Fig. 1f-h and Extended Data Fig. 4d-g. Conversely, LEVs failed to rescue KD cell defects in fibronectin deposition and organization (Fig. 1f-h and Extended Data Fig. 4h-k).

### Exosome secretion from mammary tumor-associated fibroblasts promotes *in vivo* matrix assembly, tumor growth, and metastasis

Aligned collagen fibers at the boundary of breast tumors are strongly correlated with poor patient prognosis and are thought to promote the growth and dissemination of breast cancer cells^21,22^. To test whether the *in vitro* effects of exosome secretion on stromal matrix alignment also occur *in vivo*, we injected a mixture of MDA-MB-231 breast cancer cells with scrambled control or Rab27a-KD hMFs in the mammary fat pads of NOD/SCID mice. After eight weeks, primary tumor size, lung micrometastases, and ECM characteristics were assessed. As compared to tumors containing control fibroblasts, tumors containing Rab27a-KD fibroblasts were significantly smaller and had fewer lung micrometastases (Fig. 2a-d). In addition, analysis of the stromal matrix organization within the primary tumors by immunostaining and second harmonic generation revealed that tumors containing Rab27a-KD fibroblasts had a significant reduction in aligned fibronectin, tenascin C, and collagen (Fig. 2e-g). These data are consistent with a key role for fibroblast exosomes in driving stromal matrix assembly in primary breast tumors. Since collagen that is aligned perpendicularly to the edge of breast tumors, in a “tumor-associated collagen signature-3” (TACS-3), is associated with poor patient outcome ^22^, we also analyzed second harmonic generation data from the mouse mammary tumors for this signature. Indeed, we found that tumors containing Rab27a-KD fibroblasts had a reduction in TACS-3 perpendicular fibers as compared to controls, instead having the lower-risk parallel fiber TACS-2 configuration at the periphery of tumors (Fig. 2h-j).

**Figure 2.**
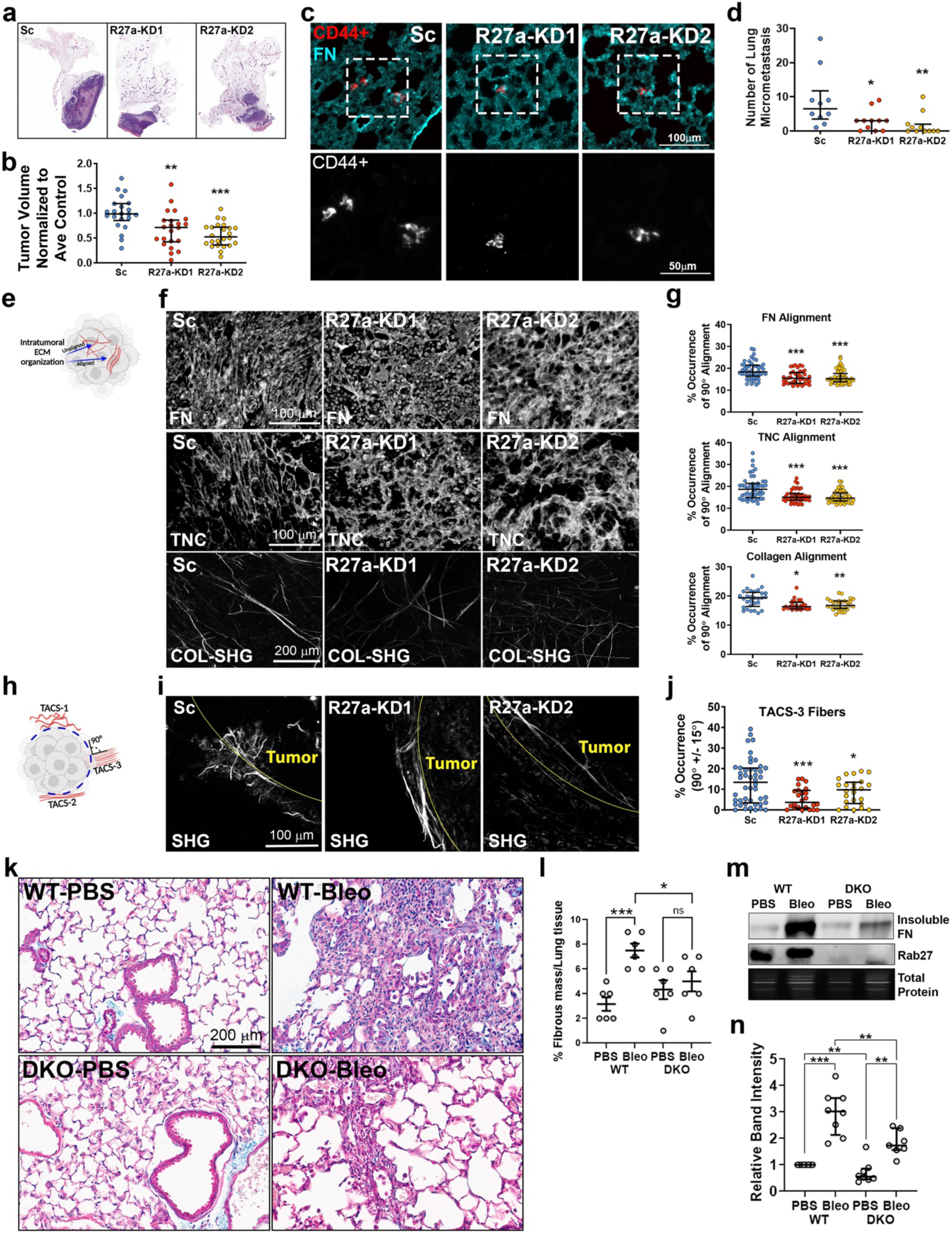
Exosome secretion promotes stromal matrix assembly and fibrosis. a, Representative H&E images of co-culture MDA-MB-231 + mammary fibroblast (scrambled control (Sc), or Rab27a-knockdown (R27a-KD1 or R27a-KD2)) mammary tumors. **b**, Tumor volume quantified as a change from the average Sc control tumor volume. N = 11 mice per cohort with 2 tumors per mouse. **, *p* < 0.01; *** *p* < 0.001 compared to average control (Sc) by Welch *t*-test. **c**, Representative immunofluorescent images (top panel) of metastatic MDA-MB-231 cells (CD44+, red) in the lung parenchyma counterstained with fibronectin (FN, cyan). Insets (bottom panel) show magnified CD44+ cells. **d,** Quantification of micrometastasis (≤ 10 CD44+ cell clusters) in lung of each cohort. The left lung lobe from each mouse was sectioned with 50 μm spacing and processed for immunofluorescence of CD44+ MDA-MB-231 cells. *, *p* < 0.05; **, *p* < 0.01 compared to Sc control. Significant differences were determined by Mann-Whitney *U* test. **e**, Schematic cartoon of aligned vs unaligned intratumoral ECM. **f**, Representative images of intratumoral FN (top panels), tenascin C (TNC) (middle panels), and collagen second harmonic generation (COL-SHG) (bottom panels) in Sc, R27a-KD1, and R27a-KD2 tumors. Images were collected with a 20x objective. **g**, The angle of ECM fiber orientation relative to the horizontal was analyzed in 6 fields of view from each section using orientation J (ImageJ/Fiji). Quantification of the percentage of fibers occurring at 90 degrees for FN, TNC, and collagen. *, *p* < 0.05; **, *p* < 0.01; *** *p* < 0.001 comparison to average Sc by Mann-Whitney *U* test. **h**, Schematic cartoon of tumor-associated collagen signatures (TACS 1-3). **i,** Representative SHG images of collagen fibers at the tumor-stromal boundary. **j**, Quantification of the percentage of TACS-3 fiber orientation (90 degrees +/-15 degrees). 6 - 8 images per tumor, N = 12 mice per cohort with 2 tumors per mouse. *, *p* < 0.05; *** *p* < 0.001 compared to average control (Sc) by Welch *t*-test. **k - n**, Analysis of lung fibrosis from WT vs Rab27a/b double knockout mice (DKO) treated with vehicle control (PBS) or bleomycin (Bleo). **k**, Representative Masson’s Trichrome-stained images of lung tissue sections. Scale bar: 200 µm. **l**, Quantification of % fibrous mass per lung tissue in the WT and DKO groups treated with PBS or Bleo. **m**, Representative Western blot images of FN and pan-Rab27 in WT and DKO groups treated with PBS vs Bleo. **n**, Quantification of relative band intensity for FN from **m**. Data are represented as mean ± SEM. Significant differences between groups (*, *p* < 0.05; **, *p* < 0.01; ***, *p* < 0.001).

### Exosome secretion promotes fibrosis in bleomycin-injured lungs

Fibronectin assembly also contributes heavily to pulmonary fibrosis, which can be modeled in mice by lung injury with the chemotherapy drug bleomycin^23^. To test whether exosome secretion may contribute to fibrosis in this model, 6-week-old control and Rab27a/b double knockout (DKO) mice were treated with bleomycin. After 2 weeks, the mice were sacrificed and their lungs harvested for analysis. Trichrome staining analysis of lung sections for collagen revealed that the number of fibrotic lesions was increased in bleomycin-treated control lungs compared to bleomycin-treated DKO lungs (Fig. 2k and l). Extraction of lung tissue with a deoxycholate-containing buffer that leaves behind insoluble, assembled ECM^24^, followed by SDS extraction and Western blot analysis, revealed a significant increase in insoluble fibronectin in control-treated lungs compared to DKO bleomycin-treated lungs (Fig. 2m and n).

### Purified SEVs induce fibronectin assembly in a cell-free purified component system

To test whether exosomes can directly induce fibronectin assembly, we developed a novel *in vitro* purified components system (Extended Data Fig. 5a). In this assay, cellular fibronectin purified from hMF conditioned media (Extended Data Fig 5b) and either LEVs or cushion density gradient-purified SEVs are mixed in a PCR tube and incubated for 72h at 37°C in the presence of Ca^++^ and Mg^++^ to allow integrin activation (see methods). The mixture is then incubated with coverslips overnight at 4°C before fixation and immunostaining for fibronectin and EV markers (TSG101, CD63, or HSP70). Imaging was performed by either wide-field or confocal microscopy, as indicated in the legends. Cellular fibronectin, LEVs, or SEVs alone showed minimal staining for fibronectin structures (Fig. 3a and c, Extended Data Fig. 5c). By contrast, co-incubation of fibronectin and either LEVs or SEVs led to the appearance of fibrillar fibronectin structures. Notably, an equal number of SEVs was much more potent in inducing fibronectin structures compared with LEVs (Fig. 3a and c). We also tested whether SEVs induce fibrillar structure formation by plasma fibronectin and found that it was ineffective in this assay, suggesting a requirement for unique domains found in cellular fibronectin (Extended Data Fig. 5d) ^25,26^.

**Figure 3.**
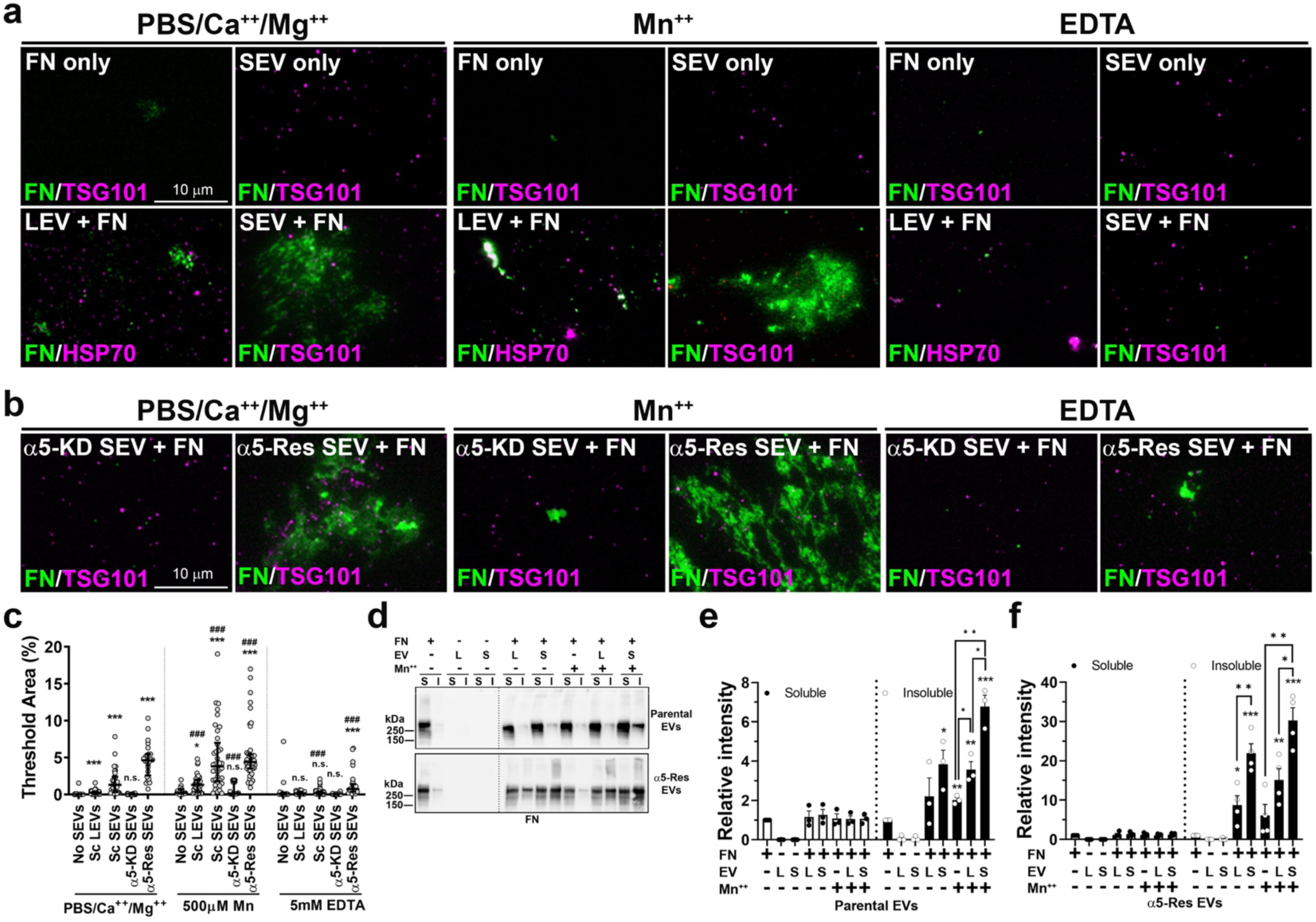
Purified SEVs can induce fibronectin assembly. **a** and **b,** Representative wide-field images for cell-free purified component assay for fibronectin assembly by parental EVs (**a**) and integrin α5-manipulated EVs (**b**) immunostained with fibronectin (FN) and HSP70 for LEVs or TSG101 for SEVs in the presence of 500 μM MnCl_2_ or 5 mM EDTA in PBS/Ca^++^/Mg^++^. α5-KD, knockdown expression of integrin α5. α5-Res, rescued expression of integrin α5. **c,** Quantitation of threshold area for fibronectin assembly (median with interquartile range) from **a** and **b**. n.s., not significant; *, *p* < 0.05; ***, *p* < 0.001 compared to No SEVs (FN only) in each condition. ###, *p* < 0.001 compared to the same EVs in PBS/Ca^++^/Mg^++^ condition. n ≥ 16 images from 3 independent experiments. Significant differences were determined by Mann-Whitney *U* test. **d,** Representative Western blots of deoxycholate (DOC) solubility assay. L, large EVs. S, small EVs. S, soluble. I, insoluble. **e** and **f,** Quantitation of parental EVs (**e**) and integrin α5-Res EVs (**f**) (mean ± SEM) from **d**. *, *p* < 0.05; **, *p* < 0.01 ***, *p* < 0.001 compared to FN only unless there is a comparison bracket. Significant differences were determined by two-tailed Student’s *t*-test. n = 4.

To test the role of integrin EV cargoes in this process, we carried out the reaction in the presence of manganese (Mn^++^) to activate integrins or EDTA to chelate divalent cations and inhibit integrins. Indeed, Mn^++^ greatly enhanced, whereas EDTA greatly reduced EV-induced fibril formation (Fig. 3a and c). As SEVs had the most activity in our assay and the fibronectin receptor α5β1 is present on SEVs (Extended Data Fig 5e) and known to be important in cellular fibronectin assembly^27^, we tested whether α5 integrin is important for SEV-induced fibronectin assembly. Indeed, many fewer fibronectin fibrils were present when the assay was carried out using SEVs purified from integrin α5-knockdown fibroblasts (Fig. 3b and c). SEVs purified from α5-KD cells in which integrin α5 was re-expressed (α5-Res) had even more robust activity in this assay, potentially due to the 3-fold increase in α5 expression over control fibroblasts (Fig. 3b and c, Extended Data Fig. 5f).

To determine whether the apparent fibrils in our images really represent assembled fibronectin, we extracted reactions from the purified components assay with 2% deoxycholate, which is known to solubilize non-assembled fibronectin^28^. Insoluble assembled fibronectin was recovered by centrifugation of the reactions at 15,000 x *g*. Western blot analysis of the supernatant and pellet fractions revealed that co-incubation of purified EVs with cellular fibronectin led to the presence of insoluble fibronectin, whereas little was detectable in the reaction that did not contain EVs. Similar to the microscopy assay, SEVs were much more potent than LEVs in inducing fibronectin assembly, and all assembly was enhanced in the presence of Mn^++^ (Fig. 3d and e). In addition, if the reactions were carried out with the α5-Res EVs that have increased α5 integrin (Extended Data Fig. 5f), EV-induced fibronectin assembly was enhanced (compare Fig. 3e to Fig. 3f, note Y-axis scale bars).

To visualize the structure of the insoluble fibronectin associated with SEVs, we carried out scanning electron microscopy of our assays, using colloidal gold-labeled fibronectin^29^. Consistent with our light microscopy and biochemical analyses, there were very few fibrillar structures associated with 72 h reactions containing only purified cellular fibronectin or only purified SEVs (Fig. 4). By contrast, reactions containing both SEVs and cellular fibronectin contained numerous gold-labeled fibrils in fibrillar arrays around the EVs. However, reactions with plasma fibronectin did not show the same fibrils but rather had many fewer and short fragments of fibrils. Reactions of SEVs and cellular fibronectin in the presence of Mn^++^ greatly enhanced fibril assembly such that the EVs were no longer visible but buried within the fibrils. By contrast, exposure of fibronectin alone to Mn^++^ did not induce fibrillogenesis by EVs+cellular fibronectin (Fig. 4). As in the light microscopy assay (Extended Data Fig. 5d), reactions with plasma fibronectin and SEVs did not show the same extensive fibrillar network as reactions with cellular fibronectin, with only a few apparent strands bound to the EVs, both in the presence and absence of Mn^++^.

**Figure 4.**
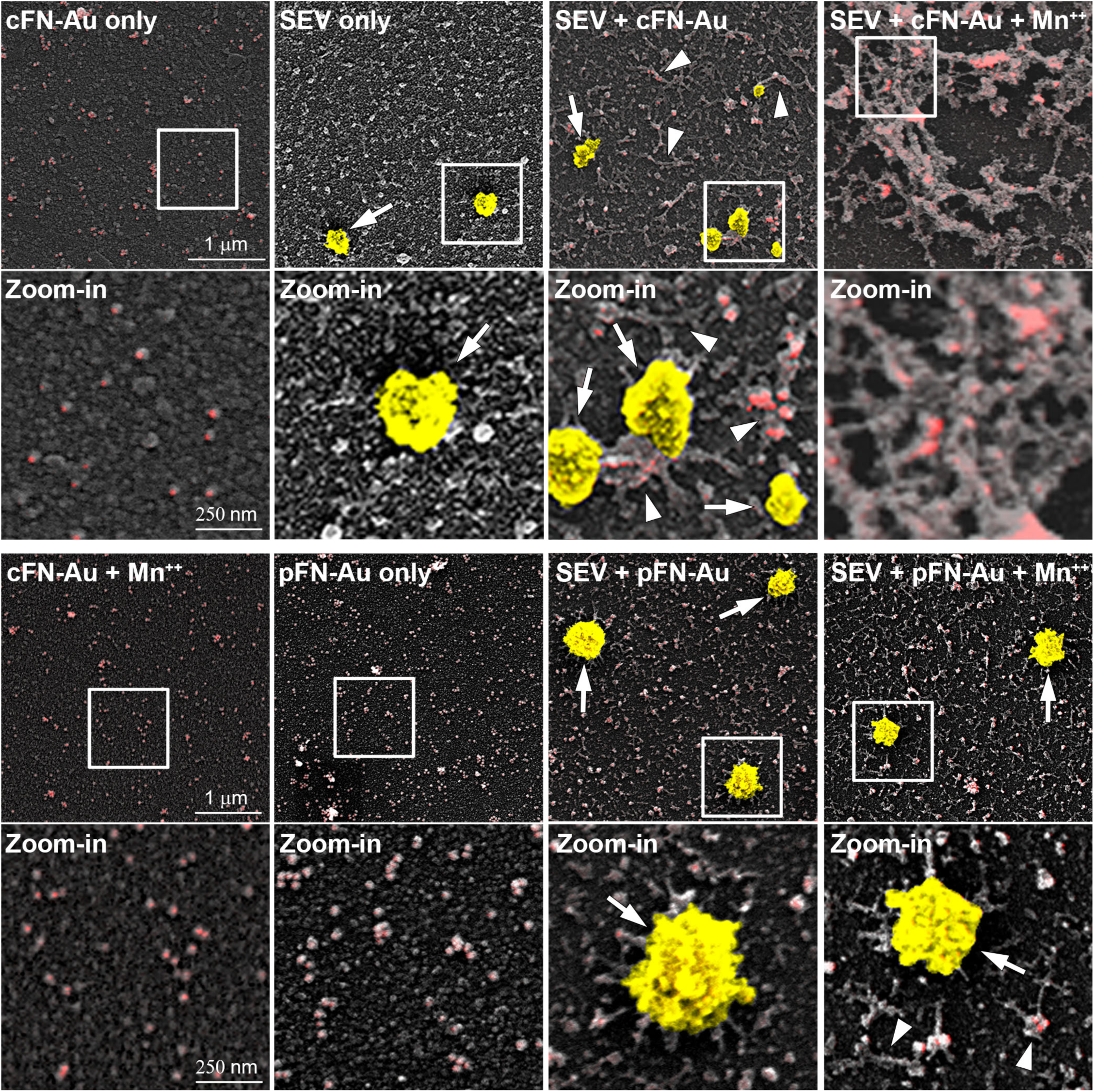
Cellular fibronectin, not plasma fibronectin, is assembled into fibrils by SEVs in cell-free purified component assay. Representative scanning electron micrographs of gold-labeled fibronectin assembly. The images in white squares are enlarged below. Arrows indicate SEVs in yellow. Arrow heads indicate FN fibrils and are omitted in SEV + cFN-Au + Mn^++^. Gold-labeled FN (FN-Au) is shown in red. cFN, cellular FN. pFN, plasma FN.

### Multiple integrins are enriched in small EVs and promote fibronectin assembly

Since SEVs are more potent than LEVs at inducing fibronectin assembly, we hypothesized that SEVs are enriched for assembly-driving cargoes. To identify such molecules, we performed quantitative proteomics using tandem mass tagging (TMT), comparing protein cargoes in SEVs with those in LEVs. As expected, SEVs were enriched (>2-fold) in tetraspanins and SEV biogenesis molecules (e.g., Alix/PDCD6IP, TSG101, Rab35), compared with LEVs (Extended Data Fig. 6a, Supplementary Data 1). While all integrins were enriched in SEVs compared to LEVs, some integrins were predicted to be enriched >2-fold in SEVs (α4, β1, β5, Extended Data Fig. 6a and b). Functional enrichment analysis showed that genes enriched in SEVs are involved in integrin-related interactions, receptor signaling networks, and syndecan-1-mediated signaling (Extended Data Fig. 6c). Additionally, a STRING analysis of protein-protein interactions indicated strong connections among integrin-ECM-related proteins enriched in SEVs (Extended Data Fig. 6d). Western blot analysis confirmed enrichment of multiple integrins in SEVs compared to LEVs (Fig. 5a).

**Figure 5.**
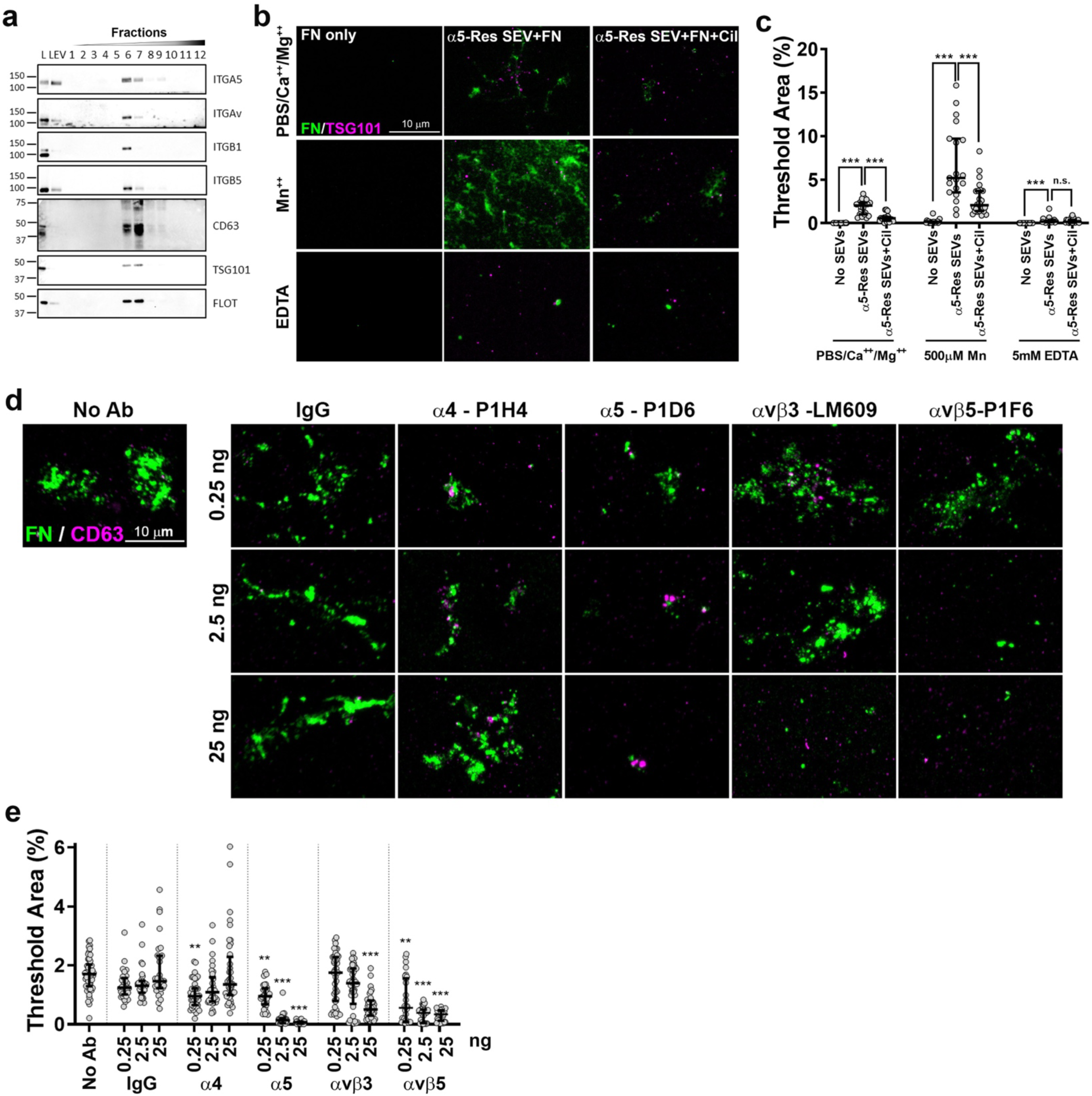
Integrins abundant in SEVs promote fibronectin assembly. a, Western blot analysis of the density gradient fractions of EVs for integrins. L, cell lysate. LEV, large EV. **b,** Representative wide-field images for cell-free purified component assay for fibronectin assembly by α5-Res SEVs with Cilengitide (Cil) immunostained with fibronectin (FN) and TG101 in the presence of 500 μM MnCl_2_ or 5 mM EDTA in PBS/Ca^++^/Mg^++^. α5-Res, rescued expression of integrin α5. **c,** Quantitation of threshold area for fibronectin assembly (median with interquartile range) from **b**. n.s., not significant; ***, *p* < 0.001. n ≥ 20 images per condition from 3 independent experiments. Significant differences were determined by Mann-Whitney *U* test. **d,** Representative wide-field images for cell-free purified component assay for fibronectin assembly by parental SEVs with integrin blocking antibodies immunostained with fibronectin (FN) and CD63 in PBS/Ca^++^/Mg^++^. IgG, immunoglobulin G. **e,** Quantitation of threshold area for fibronectin assembly (median with interquartile range) from **d**. **, *p* < 0.01; ***, *p* < 0.001 compared to the same amount of IgG. n ≥ 30 images per condition from 3 to 5 independent experiments. Significant differences were determined by Mann-Whitney *U* test.

To further investigate whether integrins enriched in SEVs contribute to SEV-induced fibronectin assembly, we first tested the effect of cilengitide, a cyclic arginine-glycine-aspartic acid (RGD)-containing pentapeptide, in the purified component assay with integrin α5-rescued SEVs. Cilengitide has the strongest activity against αvβ3 and αvβ5 integrins, with 10-fold lower activity against α5β1^30^. Cilengitide reduced fibronectin assembly even with Mn^++^ (Fig. 5b and c), suggesting that αv integrins may cooperate with α5 integrin in EV-mediated fibronectin assembly.

To test which specific SEV-enriched fibronectin-binding integrins (Extended Fig. 6b) drive fibronectin assembly, integrin-blocking antibodies were added to the assay. Immunoglobulin G (IgG) as a negative control did not inhibit fibronectin assembly, regardless of the amount (Fig. 5d and e). However, blocking antibodies for integrins α5, αvβ3, and αvβ5, (P1D6, LM609, and P1F6, respectively)^31–33^, all inhibited fibronectin assembly by SEVs in the purified component assay (Fig. 5d and e). Interestingly, the blocking antibodies to α5 and αvβ5 were effective at all concentrations tested, whereas the blocking antibody to αvβ3 was only effective at the highest concentration tested. By contrast, a blocking antibody to integrin α4 (P1H4)^34^ had little effect on cellular fibronectin assembly, despite its strong enrichment in SEVs, with some inhibition at the lowest antibody concentrations but no dose-dependency (Fig. 5d and e, Extended Fig. 6b). These data indicate that α5β1, αvβ3, and αvβ5 are all key SEV cargoes that induce fibronectin assembly.

**Figure 6.**
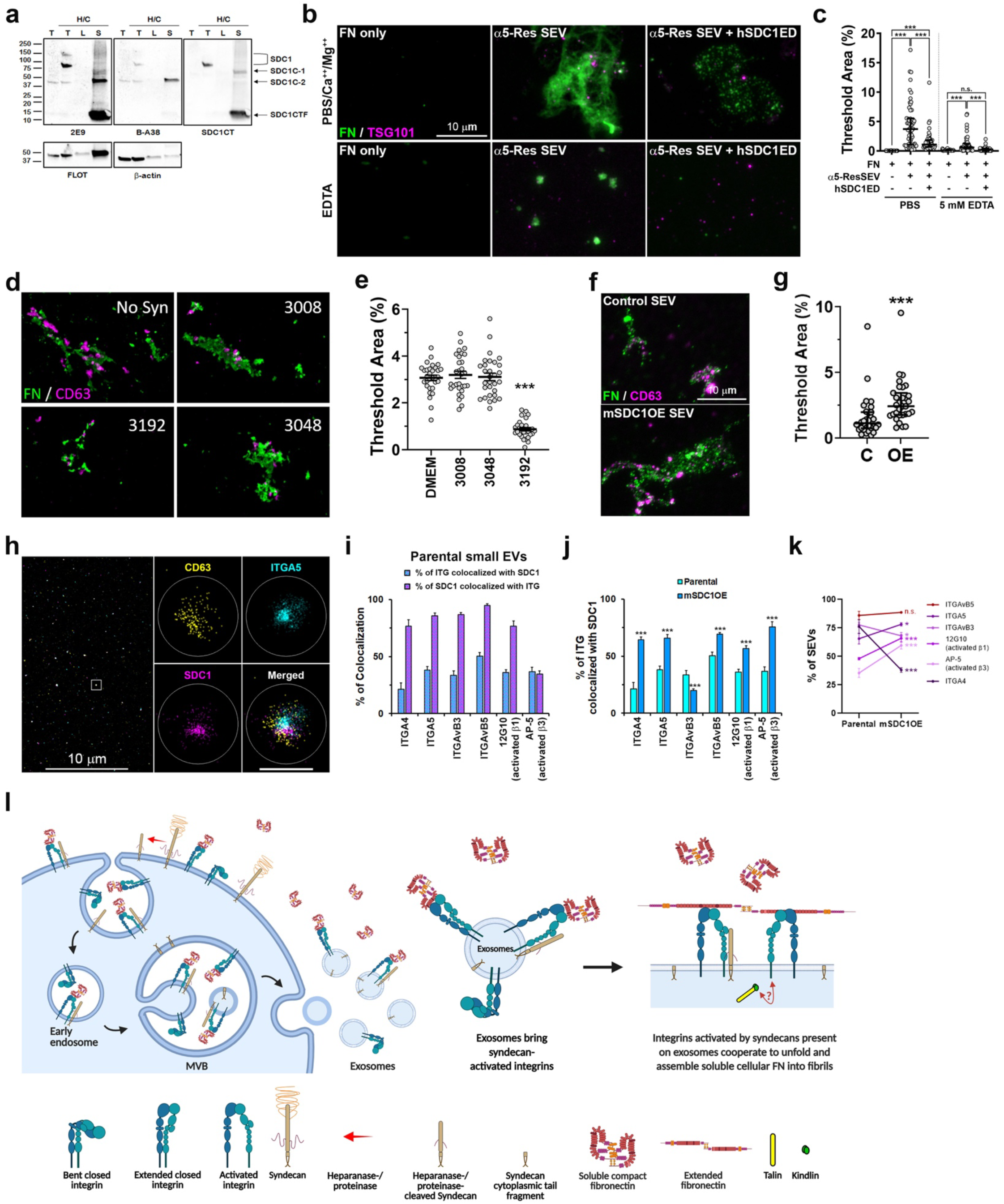
Integrins and syndecan synergize in the same SEVs to promote fibronectin assembly. a, Western blot analysis for syndecan 1 (SDC1) treated with heparinase/chondroitinase (H/C). T, total cell lysate. L, large EV. S, small EV. SDC1C, syndecan 1 cleaved form. SDC1CTF, syndecan 1 cytoplasmic tail fragment. **b,** Representative wide-field images for cell-free purified component assay for fibronectin assembly by α5-Res SEVs with recombinant human syndecan 1 ectodomain (hSDC1ED) immunostained with fibronectin (FN) and TG101 in the presence of 5 mM EDTA in PBS/Ca^++^/Mg^++^. α5-Res, rescued expression of integrin α5. **c,** Quantitation of threshold area for fibronectin assembly (median with interquartile range) from **b**. n.s., not significant; ***, *p* < 0.001. n ≥ 39 images from 3 or 4 independent experiments. **d,** Representative wide-field images for cell-free purified component assay for fibronectin assembly by α5-Res SEVs with synstatin peptides immunostained with fibronectin (FN) and CD63 in PBS/Ca^++^/Mg^++^. α5-Res, rescued expression of integrin α5. **e,** Quantitation of threshold area for fibronectin assembly (median with interquartile range) from **d**. ***, *p* < 0.001 compared to control (No Syn). No Syn, DMEM with no synstatin. n = 30 images from 3 independent experiments. **f,** Representative wide-field images for cell-free purified component assay for fibronectin assembly by mSDC1OE SEVs (mouse syndecan 1-overexpressed small EVs) immunostained with fibronectin (FN) and CD63 in PBS/Ca^++^/Mg^++^. **g,** Quantitation of threshold area for fibronectin assembly (median with interquartile range) from **f**. ***, *p* < 0.001 compared to control SEVs (C). OE, mouse syndecan 1-overexpressed SEVs. n = 30 images from 3 independent experiments. **h,** Representative super-resolution microscopy images of small EVs immunostained for integrin α5, SDC1, and CD63. The image in a white square is enlarged at the right. Scale bar in an enlarged image, 200 nm. **i,** Percentage of colocalization of integrins and SDC1 in the same small EVs. **j,** Comparison of the percentage of integrins colocalized with SDC1 in the same small EVs purified from parental and mouse SDC1-overexpressing hMFs represented as mean ± SEM. ***, *p* < 0.001 compared to parental SEVs. n ≥ 15 images from 3 or 4 independent experiments. Significant differences were determined by unpaired *t*-test. **k,** Percentage of integrin-containing small EVs represented as mean ± SEM. n.s., not significant; *, *p* < 0.05; ***, *p* < 0.001 compared to parental SEVs. n ≥ 15 images from 3 or 4 independent experiments. Significant differences were determined by unpaired *t*-test. **l,** Proposed model for FN fibril assembly by exosomes. Processing of syndecans by heparanase and proteinase at the cell surface promotes endocytosis and biogenesis of exosomes ^65,66^. Likewise, activated integrins are endocytosed and sorted to MVBs ^67^, possibly in association with processed forms of syndecan-1. Exosomal integrins bind soluble cellular FN, and the bound FN can be unfolded and interact with other integrins. Clustering of activated integrins, either within or from neighboring exosomes, further stretches cellular FN into fibrillar form. Syndecan 1 is mainly present in exosomes as a cleaved form, which often contains a peptide region important for binding and activating integrins. Talin-1, kindlin-2/FERMT2, and syntenin-1 present in the exosomal lumen (Supplementary Data 2, Extended Fig. 7e) may also promote integrin activation and clustering.

### Syndecan-1 synergizes with integrins to promote fibronectin assembly

In addition to integrins, other fibronectin- and integrin-binding molecules were enriched in SEVs (Supplementary Data 1, Extended Fig. 6b-d). Of these, the exosome biogenesis molecule syndecan-1 was chosen as a likely candidate to synergize with integrins in driving exosomal fibronectin assembly, based on its known activity in driving cellular fibronectin assembly and organization, its integrin binding activities, and its strong enrichment in fibroblast SEVs^35,36^ (Extended Fig. 6b, Fig. 6a). 2E9^37^ and SDC1CT (D4Y7H, Cell Signaling Technology) antibodies bind to peptide epitopes in the cytoplasmic tail, and B-A38 antibody recognizes the ectodomain of syndecan-1. Consistent with a previous report^38^, we found that the major form of syndecan-1 in SEVs was a cleaved C-terminal fragment. However, we also identified several ectodomain fragments of syndecan-1 that might interact with integrins and/or fibronectin (Fig. 6a, Extended Fig. 7a).

To investigate whether syndecan-1 promotes fibronectin assembly by SEVs, purified recombinant syndecan-1 ectodomain was added to the cell-free purified component reactions. These ectodomains bind to integrins and other syndecan-binding transmembrane proteins, but not fibronectin, as they are produced in bacteria and lack the heparan sulfate groups. While a small amount of fibronectin structure formation was observed, it was greatly reduced compared to control reactions containing only both SEVs and fibronectin (Fig. 6b and c). As a second inhibitory approach, synthetic peptides, called synstatins, corresponding to syndecan domains that interact with other proteins^39^, were added to the purified components assay. Interestingly, whereas peptides targeting the interaction of syndecan-1 (3008) or syndecan-4 (3048) with integrin α3β1 had no effect, “synstatin_93-120_,” a peptide that competitively blocks the interaction of syndecan-1 with integrins αvβ3 and αvβ5^40^ inhibited fibronectin assembly (3192) (Fig. 6d and e). Notably, the peptide that corresponds to synstatin (aa93-aa120) overlaps with one of the 4 peptides identified by mass spectrometry to be present in SEV-carried syndecan-1 (Extended Fig. 7a, aa101-aa117 underlined and bolded in red), providing evidence that ectodomain-including syndecan-1 is present in fibroblast SEVs and is important for fibronectin assembly.

To further test whether syndecan-1 can drive SEV-mediated fibronectin assembly, we overexpressed (OE) mouse syndecan-1 in hMFs (Extended Fig. 7b). Purified control and OE SEVs were tested in the purified components assay. Indeed, syndecan-1 OE SEVs induced significantly more fibronectin fibrillar structures than control SEVs (Fig. 6f and g).

To determine whether syndecan-1 is present in the same small EVs as integrins, mouse syndecan-1 was tagged with mScarlet protein, expressed in hMFs, and then immunoprecipitated from the lysates of purified small EVs using RFP-Trap magnetic beads (Extended Fig. 7c). The immunoprecipitated complex was analyzed by mass spectrometry, and three peptides were identified from the ectodomain of mouse syndecan-1 (Extended Fig 7d).

Further STRING analysis with co-precipitated proteins identified by the mass spectrometry revealed integrins and other proteins involved in extracellular matrix organization, integrin-mediated signaling, and focal adhesion assembly. Notably, proteins involved in integrin activation, e.g., talin-1 (TLN1) and kindlin-2 (FERMT2), were present in the immunoprecipitate (Extended Fig. 7e).

As a second approach to determine whether syndecan-1 is present with integrins in the same EVs, small EVs were immunostained with antibodies against syndecan-1, various integrins, and the EV marker CD63 in three-color reactions, and analyzed by direct stochastic optical reconstruction microscopy (dSTORM) (Fig. 6h). Indeed, we found that integrins were frequently in the same EVs as syndecan-1. Furthermore, we could detect activated integrins in small EVs (Extended Fig 8a and Fig 6i). In control hMF EVs, generally more than 75% of syndecan-1 containing SEVs also contained integrins (Fig. 6i, purple bars). Conversely, a lower percentage of integrin-containing EVs also contained syndecan-1, suggesting that integrins are a more prevalent cargo under baseline conditions (Fig 6i). Indeed, in EVs purified from syndecan-1-overexpressing hMFs, the percentage of integrin-positive EVs that also had syndecan-1 generally increased (Fig. 6j). Most notably, the percentage of EVs that were positive for activated integrins β1 and β3 increased (Extended Fig. 8a,b and Fig 6j,k), suggesting that syndecan-1 in EVs promotes integrin activation, which is important for fibronectin assembly. As dSTORM is based on blinking fluorophores, we were able to determine the number of molecular localizations per EV in our images. Analysis of the integrin-positive EVs for molecular number revealed that overexpression of syndecan-1 led to an increase in the number of integrin αvβ5 molecules as well as of activated β1 molecules per EV. Thus, syndecan-1 increases both the percent of EVs containing integrins, as well as their number and activation status, especially of the integrins involved in fibronectin assembly (Fig. 6j, Extended Fig. 8b,c, Fig. 5d,e).

## Discussion

Our study challenges the longstanding paradigm that fibronectin matrix assembly is exclusively a cell-mediated process by identifying exosomes as autonomous agents capable of initiating fibronectin fibrillogenesis. Traditionally, fibronectin fibril formation has been understood as requiring direct cell engagement via integrins and cytoskeletal tension ^1,41^. In contrast, we demonstrate that exosomes, independent of intact cells, can drive the assembly of fibronectin fibers, revealing a previously unappreciated extracellular mechanism of matrix remodeling.

This discovery fundamentally expands the functional repertoire of extracellular vesicles (EVs), positioning exosomes not merely as carriers of cellular signals^42^ but as active architects of the extracellular matrix. Our data show that exosome surfaces provide a platform that nucleates fibronectin fibrillogenesis, leveraging integrin and syndecan interactions reminiscent of those on the cell surface ^1,41^, but likely more potent due to the concentration in EVs of active integrins and syndecan-1, which is in a processed form distinct from that on the cell surface (Fig. 6l). This ability of exosomes to organize and remodel the ECM implicates them as key mediators of microenvironmental remodeling, with profound implications for diseases characterized by pathological matrix accumulation.

Fibrosis and cancer, both marked by excessive ECM deposition and altered matrix architecture, may be directly influenced by exosome-mediated fibronectin assembly, as we show for bleomycin-induced lung fibrosis and breast tumor-associated stromal matrix assembly. The pro-fibrotic microenvironment, historically attributed to aberrant fibroblast activation, could also arise from increased exosome production or altered EV cargoes (such as syndecan-1 ^43–45^) that promote fibrillogenesis extracellularly. Similarly, in cancer, exosomes could facilitate metastatic progression not only through cargo delivery but by restructuring the ECM to create aligned collagen and fibronectin fibers that guide invasive cells ^46^.

This paradigm shift underscores the need to reconsider EVs as dynamic extracellular effectors in tissue remodeling rather than passive messengers. The syndecan-syntenin-ALIX axis, previously linked to exosome biogenesis ^38^, may couple matrix assembly with vesicle production, coordinating EV secretion and ECM organization in a feed-forward loop.

Therapeutic interventions targeting exosome production, release, or surface composition could thus offer novel strategies to disrupt pathological ECM remodeling in fibrosis and metastatic cancer.

Overall, our findings uncover exosomes as critical extracellular organizers of fibronectin matrices, redefining their role in the tumor microenvironment and fibrotic niches. By discovering this unexpected mechanism of matrix assembly, we open new avenues for understanding and ultimately intervening in diseases driven by aberrant ECM deposition.

## Methods

### Cell culture and reagents

Human primary mammary fibroblasts (hMFs) obtained after reduction mammoplasty and immortalized using hTERT (generous gift from Kuperwasser lab, Tufts University) were maintained in high glucose Dulbecco’s Modified Eagle Medium (DMEM) supplemented with 10% bovine calf serum (CS) at 37°C with 5% CO_2_. For EV isolation, cells were cultured in serum-free conditions using Opti-MEM for 48h. Human prostatic cancer-associated fibroblasts (CAFs) were isolated from prostate cancer tissue and maintained in RPMI-1640 supplemented with 10% fetal bovine serum (FBS) at 37°C with 5% CO_2_ (generous gift from Dr. Donna Webb, our late colleague). Mouse cardiac stromal cells derived from immortomice were obtained from Dr. Igor Feoktistov (Vanderbilt University Medical Center) and maintained on 0.1% gelatin-coated tissue culture plates in DMEM supplemented with 10% fetal bovine serum, 2 mM glutamine, and 10 ng/ml IFNγ at 33°C with 5% CO_2_. Cells were replated and cultured without IFNγ at 37°C with 5% CO_2_, 6 days before experiments^47^. MDA-MB-231 cells were obtained from American Type Cell Culture (ATCC) and maintained in DMEM supplemented with 10% fetal bovine serum at 37°C with 5% CO_2._ A lentiviral shRNA expression system, pLKO.1, was used to knock down Rab27a (Thermo Fisher Scientific TRCN0000005296 (5’-CCGG-CGGATCAGTTAAGTGAAGAAA-CTCGAG-TTTCTTCACTTAACTGATCCG-TTTTT-3’) and TRCN0000005297 (5’-CCGG-GCTGCCAATGGGACAAACATA-CTCGAG-TATGTTTGTCCCATTGGCAGC-TTTTT-3’)), synaptotagmin-7 (Open Biosystems RHS3979-9577052 and RHS3979-9577053), integrin α5 3’ UTR (Sigma-Aldrich TRCN0000230126 (5’-AGGCAGATCCAGGACTATATT-3’) ) or scrambled control (Addgene #26701 (5’-CCTAAGGTTAAGTCGCCCTCG-3’)). Human integrin α5 construct (pEGFP-N3-ITGA5) was obtained from Dr. Webb. Integrin α5-GFP was cloned into a pLenti6/V5-DEST vector using the Gateway Vector Kit (Invitrogen). The GFP was replaced with mScarlet from pCytERM-mScarlet-N1 (Addgene #85066) using the Gibson Assembly Master Mix (NEB E2611) with primer sets (mScarlet forward, 5’-TCCATCGCCACCATCATGGTGAGCAAGGGC-3’; mScarlet reverse. 5’-TCGGCGCGCCCACCCTTCTAGGATCCCTTGTACAGCTCGT-3’; pLenti vector forward, 5’-ACGAGCTGTACAAGGGATCCTAGAAGGGTGGGCGCGC-3’; pLenti vector reverse, 5’-CCCTTGCTCACCATGATGGTGGCGATGGATCC-3’). pLenti6/V5-DEST-ITGA5-mScarlet was used to rescue the expression of integrin α5 in integrin α5-knockdown hMFs. Full-length mouse syndecan-1 cDNA (pcDNA3; provided by R. Sanderson, University of Alabama at Birmingham) was cloned into a pLenti CMV vector using Gibson Assembly Master Mix with primer sets (mSDC1 forward, 5’-ACAAAAAAGCAGGCTCCACCATGAGACGCGCGGCGC-3’; mSDC1 reverse, 5’-CAAGAAAGCTGGGTCTAGAGGCGTAGAACTCCTCCTGC-3’; pLenti vector forward, 5’-CTAGACCCAGCTTTCTTGTACAAAG-3’; pLenti vector reverse, 5’- AGCCTGCTTTTTTGTACAAACTTGT-3’). To create an mSDC1-mScarlet construct, two complementary strands of a (GGGGS)3 linker containing BamH I and Xba I sites at the 3’ end were synthesized first (5’- GGAGGAGTTCTACGCCTCAGGTGGAGGCGGTTCAGGCGGAGGTGGCTCTGGCGGTGGCGGATCCAAAAATAAAAAATCTAGACCCAGCTTTCTTG-3’). The complementary strands were annealed to each other and cloned at the C-terminus of mSDC1 using NEBuilder HiFi DNA Assembly (NEB E5520). mScarlet was amplified from pCytERM-mScarlet-N1 by adding BamH I and Xba I sites at the 5’ and 3’ ends, respectively (forward primer, 5’- CGCGGATCCGTGAGCAAGGGC-3’; reverse primer, 5’- TGCTCTAGAGCATTACTTGTACAGCTCG-3’). The amplified mScarlet fragment and pLenti CMV-mSDC1-linker were digested by BamH I and Xba I. pLenti CMV-mSDC1-linker-mScarlet was used to produce hMFs stably expressing mSDC1-mScarlet. Viral particles were produced from 283FT cells by cotransfection with viral vectors. Cells were transduced by viral particles and selected using selection markers. Primary antibodies were: anti-CD63 (abcam ab68418, 1:500 for WB and ab8219, 1:100 for IF), anti-TSG101 (abcam ab30871, 1:1,000 for WB and 1:100 for IF), anti-flotillin (BD Biosciences 610820, 1:1,000 for WB), anti-HSP70 (sc-373867, Santa Cruz Biotechnology, 1:100 for IF), anti-GM130 (BD Biosciences 610822, 1:250 for WB), anti-Rab27a (Cell Signaling 69295, 1:1,000 for WB), anti-pan-Rab27 (OriGENE SKU AB7223, 1:500 for WB), anti-synaptotagmin-7 (Synaptic System 105173, 1:200 for WB), anti-fibronectin (Sigma-Aldrich F3648, 1:3,000 for WB and 1:300 for IF; BD Bioscience 610077, 1:5,000 for WB and 1:500 for IF; abcam ab23750, 1:500 for IF on tissues), anti-tenascin C (Millipore MAB1911, 1:500 for WB and IF; abcam ab108930, 1:1,000 for IF on tissues), anti-CD44 (abcam ab51037, 1:2,000 for IF on tissues), anti-integrin α4 (Cell Signaling 8440 (D2E1), 1:5,000 for WB), anti- integrin α5 (Santa Cruz Biotechnology 166665 (clone A11), 1:3,000 for WB), anti-integrin αv (abcam 208012, 1:1,000 for WB), anti-integrin β1 (BD Biosciences 610467 (clone 18), 1:1,000 for WB), anti-integrin β3 (Millipore MAB1974, 1:1,000 for WB), anti-integrin β5 (abcam 184312, 1:1,000 for WB), anti-syndecan-1 (BioRad MCA2459 (B-A38), 1:2,000 for WB, Cell Signaling 12922 (D4Y7H), 1:2,000 for WB, and Zimmerman lab 2E9 hybridoma supernatant, 1:4 for WB), anti-mouse syndecan-1 (BioLegend 142502 (281-2), 1:1,000 for WB), anti-RFP (Chromotek 6G6, 1:2,000 for WB) and anti-β-actin (Ac-74, Sigma, 1:5,000 for WB). HRP-conjugated goat anti-mouse IgG (W4021, 1:10,000 for WB) or goat anti-rabbit IgG (W4011, 1:10,000 for WB)were purchased from Promega. Alexa Fluor 488-conjugated donkey anti-mouse IgG (A32723, 1:1,000 for IF) or donkey anti-rabbit IgG (A21206, 1:1,000 for IF), and Alexa Fluor 546- conjugated donkey anti-mouse IgG (A10036, 1:1,000 for IF) or goat anti-rabbit IgG (A11010, 1:1,000 for IF) secondary antibodies were purchased from Thermo Fisher Scientific. Colloidal gold (10 nm)-conjugated donkey anti-rabbit IgG was purchased from Electron Microscopy Sciences (25704, 1:40 for immunogold transmission electron microscopy). Pierce ECL Western Blotting Substrate and SuperSignal WestFemto Maximum Sensitivity Substrate (32106 and 34095, respectively, Thermo Fisher Scientific) were used as substrates for Western blotting. Blots were imaged using an Amersham Imager 600 (GE Healthcare Life Sciences) or iBright 1500 (Thermo Fisher Scientific) and analyzed using Fiji. The human syndecan-1 extracellular domain^48^ and mouse syndecan-4 extracellular domain^49^ fused to GST and inhibitory peptides against syndecan-1 and syndecan-4^39^ were previously described elsewhere.

### Cell-derived extracellular matrix preparations

The preparation of cell-derived matrix was described previously^50^. Briefly, cells were plated in complete media on glass coverslips at 80% confluency and allowed to attach overnight. The following day, the cells were washed multiple times using DMEM and cultured in DMEM supplemented with 1 ng/ml of TGFβ1 and 50 µg/ml of ascorbic acid. After 72 h, the cells were washed with PBS and lysed in 25 mM Tris-HCl, pH 7.4, 150 mM NaCl, 0.5% Triton-X100, 20 mM NH4OH for 5 min at 37°C. To ensure complete removal of the cells, plates were checked under the microscope. Cellular debris was carefully removed with a pipette, and the matrix was washed gently with PBS. For immunoblotting, matrix samples were lysed in equal volumes of 2x Laemmli sample buffer (0.1 M Tris-HCl, pH 6.8, 4% SDS, 20% glycerol, 0.004% bromophenol blue, 10% β-mercaptoethanol) for each cell line and boiled at 100°C for 5 min. For immunofluorescence, matrix samples were prepared on glass coverslips, fixed with 4% paraformaldehyde (PFA) in PBS for 15 min, and blocked with 5% bovine serum albumin (BSA) in PBS for 2h at room temperature. The samples were immunostained for fibronectin or tenascin-C. For general ECM protein staining, the samples were fixed with 4% PFA, washed with 50 mM NaHCO_3_, pH 8.7, and stained with Cy2 bis-Reactive Dye (GE Healthcare PA22000, 20 μg/ml in 50 mM NaHCO_3_, pH 8.7) for 30 min. And then, the labeling was stopped by adding the same volume of 400 mM Tris-HCl, pH 7.5. Images were acquired using a Nikon S Fluor 20x/0.75 NA lens with a Nikon Eclipse TE2000E microscope with a cooled charge-coupled device (CCD) camera (Hamamatsu ORCA-ER). Images were analyzed and quantified using ImageJ.

### Isolation of EVs by cushioned-density gradient

The Rafai group from UCSF developed the cushioned-density gradient ultracentrifugation method^20^. To collect conditioned media, 80% confluent cells were cultured for 48h in Opti-MEM. EVs were isolated from conditioned media by serial centrifugation at 300 x *g* for 10 min, 2,000 x *g* for 20 min, and 10,000 x *g* for 30 min (9,300 rpm in a Ti45 rotor, Beckman Coulter) at 4°C to respectively sediment live cells, dead cells and debris, and large EVs. The supernatant from the 10,000 x *g* centrifugation was concentrated to 30 ml and then layered on a 2 ml 60% iodixanol cushion (Sigma Aldrich D1556) and spun at 100,000 x g for 4h at 4°C using a SW32 rotor (Beckman Coulter). The 2 ml cushion and 1 ml of the concentrated supernatant (40% (w/v)) were collected and layered at the bottom of a discontinuous iodixanol gradient (20% (w/v),10% (w/v), and 5% (w/v) by diluted in 0.25 M sucrose, 10 mM Tris-HCl pH 7.5 from the bottom to the top of a 14 x 89 mm polyallomer tube. The gradient was spun at 100,000 x *g* for 18 h at 4°C using a SW40 rotor (Beckman Coulter). Twelve fractions were collected, diluted in PBS, and pelleted using a TLA110 rotor (Beckman Coulter) at 100,000 x g for 3h. The pelleted small EVs (fractions 6 and 7) were resuspended in PBS. EV number and size were measured by ZetaView (Particle Metrix).

### Matrix alignment assay in collagen gel

Cells were suspended at 5 *10^6^/ml in DMEM supplemented with 10% CS depleted with exosomes by centrifugation at 100,000 x *g* for 18 h at 4°C using a Ti45 rotor (Beckman Coulter). Collagen gel was prepared with 1x PBS,18 mM NaOH, 3 mg/ml rat tail type I collagen (Corning 354249). The same volume of cell suspension and collagen gel was mixed, and 20 μl of cell-collagen gel mix was plated on μ-Slide 8-well, ibiTreat (ibidi, 80826). After gelation for 1.5 h, DMEM supplemented with 10% exosome-depleted CS was added to each well. Medium containing isolated large EVs or small EVs was changed every 2 to 3 days for a week. The 3D culture was fixed using 4% PFA in PBS, washed using PBS, quenched using 150 mM glycine in PBS, and blocked using 5% BSA in PBS. Then, a general immunostaining procedure was performed. ECM staining was imaged using a 20x lens equipped on an LSM 510 confocal microscope (Zeiss). Z-stack images per each region were projected and analyzed for assembly and alignment of matrix using CT-FIRE software (http://www.loci.wisc.edu/software/ctfire), which combines the advantage of the fast discrete curvelet transform for denoising the image and enhancing the fiber edge features and the advantage of fiber extraction (FIRE) algorithm for extracting individual fibers ^51^.

### Animal models

NOD/SCID and Rab27a/b DKO mice were housed and handled in accordance with the Guide for Care and Use of Laboratory Animals in AAALAC-accredited facilities, and all procedures were approved by the University of Wisconsin–Madison Animal Care and Use Committee (protocol number M005840). NOD/SCID mice were purchased from Jackson Laboratories. Rab27a/b DKO mice (a kind gift from Dr. Miguel Seabra) were generated by crossing Rab27b KO mice with C57BL/6-Rab27a^ash/ash^ mice and subsequently crossing the heterozygous progeny as described^52^. For xenograft tumor studies, 1 x 10^6^ MDA-MB-231 cells plus 5 x 10^5^ RMFs, which were transfected with either scrambled shRNA (Sc) or shRNAs against Rab27a (R27a-1 and R27a-2), were washed and resuspended in 50 µl of sterile PBS. Cells were orthotopically injected bilaterally into the inguinal mammary fat pads of 8-week-old NOD/SCID female mice. After 8 weeks, tumor growth was measured by calipers and volume calculated (V = 0.5 x L x W2)^53^. Mice were humanely euthanized and tumor and lung tissues harvested for further analysis. Lung tissues were gravity-inflated with 10% neutral buffered formalin (NBF) to maintain structure during fixation. Lung and tumor tissue is then fixed in 10% NBF for 48 h at 4 °C and then paraffin-embedded by standard methods. For bleomycin-induced lung fibrosis, 6-week old C57BL/6 WT and Rab27a/b DKO mice were anesthetized with isoflurane before being administered a single intratracheal (IT) dose of bleomycin (6 U/kg, Hospira, Lake Forest, IL, USA) in 50 μL of DPBS or DPBS alone for vehicle controls, as we have done before^54,55^. Two weeks post-treatment, mice were euthanized, and lungs were harvested for further analysis. The left lung lobe was dissected and flash-frozen for western blot analysis, while the right lobes were gravity inflated and fixed in 10% formalin for 48 h at 4°C and processed for histological examination.

### Immunohistochemistry and Imaging

Immunofluorescence analysis of lung and xenograft tumor tissue was performed as previously described^56^ with antibodies outlined above. Briefly, tissues were formalin-fixed and prepared by standard methods for paraffin-embedding (FFPE) and sectioning. A set of FFPE sectioned tumor tissues were stained hematoxylin and eosin (H&E) by routine methods and imaged by Aperio Digital Pathology Slide Scanner system.

Another set of FFPE tissues were subject to standard deparaffinization, rehydration and antigen retrieval with citrate buffer for 20 min and blocked with 10% BSA for 30 min. Then the sections were subjected to the TSA Plus kit for tissue labeling following manufacturers’ protocols (Perkin Elmer, fluorescein NEL741001KT and Cy 3.5 NEL763001KT, Waltham, MA). Tissue was incubated in appropriate primary antibody for 1h at room temp. HRP-conjugated anti-rabbit (1:500) was added for 10 minutes followed by 10 min incubation with TSA Plus kit working solution, including the desired fluorophore. Tissues underwent the antigen retrieval step again for 20 min if the same tissue would be subjected to multiple labeling before being mounted in ProLong Gold with DAPI (Thermo Fisher Scientific P36931). Images were collected using a Nikon Eclipse TE300 inverted microscope with a Nikon Plan Fluor ELWD 20X/0.45 DIC L objective lens and an ORCA-R2 Digital CCD camera. A minimum of 5 images were collected from each tumor for fibronectin and TNC orientation analysis. 4 sections from each mouse lung at least 200 µm apart were imaged for micro-metastases using SlideBook Software (v6) and processed using FIJI software. A micro-metastasis was counted as a group of 3 or more CD44+ cells. FFPE lung tissue samples from the bleomycin-induced pulmonary fibrosis model were sectioned into 5 μm slices and stained with Masson’s Trichrome using standard protocols.

Slides were scanned at 20x magnification with an Aperio Digital Pathology Slide Scanner. Thirty regions of interest (ROIs) were selected for analysis using Aperio ImageScope software (v12.4.6.5003). The images were analyzed for fibrosis using Orbit Image Analysis software (Actelion Pharmaceuticals Ltd, Allschwil, Switzerland) with a previously developed training set^57^. Briefly, this method is based on the two main parameters described in the modified Ashcroft numerical scale: thickening of alveolar walls by collagen deposition and fibrotic masses.

Collagen within and surrounding a 300 μm vibratome section of primary xenograft tumors was imaged using Ultima IV 2-photon microscopy on a Nikon Eclipse Ti base (Bruker Nano Surfaces, Middleton, WI). Laser excitation on the Ultima IV was provided using a Coherent Chameleon laser with Hamamatsu multi-alkali photomultiplier detectors. Data acquisition and scanning control was done using PrairieView (Bruker Nano Surfaces, Middleton, WI). For intratumoral collagen, images were collected using a Nikon 40x 1.15 NA water immersion objective lens. For TACS analysis, 5 fields of view were collected along the tumor stromal boundary using a 20x Ziess 1.0 NA water immersion objective lens. Using 890 nm excitation, collagen second-harmonic generation emission was observed at 445 nm and discriminated from fluorescence using a 445 nm 20 nm narrow band-pass emission filter (Semrock). For each image, z-stacks of 15 planes, 3 µm apart, were projected into one image with a minimum of 5 images per tumor.

The images of intratumoral collagen in primary tumors were analyzed by OrientationJ, a plugin for FIJI, to determine the general orientation of the extracellular matrix being analyzed. This gave us the occurrence of fibers being organized in a certain degree of orientation from 0-180 degrees. These orientations were then binned every 20 degrees, and the bin in which the largest percentage of fibers was located was set to be oriented at 80-100 degrees in order to be comparable to all other images. The percentage of fibers oriented at 80-100 degrees out of the total number was averaged for all of the mice between each condition, and these averages were then analyzed for statistical significance using Prism 8.0. Collagen fiber orientation at the tumor stromal boundary (TACS) was analyzed by CT-Fire. All collagen fibers with angles of 90° +/- 15° from the tumor boundary were classified as TACS-3^21,58^.

### Western blotting with tissue samples

Flash-frozen lung tissues were pulverized using a mortar and pestle under liquid nitrogen for efficient ECM extraction. For protein extraction, the pulverized tissue was subjected to a two-step lysis protocol, as previously described^24^, separating detergent-soluble and insoluble fractions. In short, pulverized lung tissue was homogenized in RIPA buffer containing 1% sodium deoxycholate (DOC) at a concentration of 0.1 g tissue per mL buffer. After centrifugation at 4°C, the supernatant (soluble fraction) was collected, while the pellets were solubilized in a buffer containing 4 M urea, 4% SDS, and 1 mM DTT. The resuspended pellets (insoluble fraction, containing assembled ECM) were vortexed, sonicated, and heated to 95°C for 5 min to ensure complete protein solubilization. Protein concentrations were determined using the DC Protein Assay kit (Bio-Rad) with BSA standards. Proteins from the insoluble fractions were separated by SDS-PAGE and transferred to a PVDF membrane. To confirm consistent loading and transfer efficiency, the membrane was stained with Fast Green FCF (MilliporeSigma) for total protein staining. The membrane was blocked with 5% non-fat dry milk in TBS-T and incubated overnight at 4°C with primary antibodies against FN and pan-Rab27. After incubation with HRP-conjugated secondary antibodies (Jackson ImmunoResearch) and chemiluminescence substrate (LI-COR BioScience), or with fluorescent-conjugated secondary antibodies (IRDye from LI-COR Biosciences), blots were imaged using the Odyssey Fc Imager. Band intensities were quantified using Image Studio software (LI-COR Biosciences).

### Transmission electron microscopy

Formvar carbon film-coated grids were rinsed by immersion into double-distilled water and 95% ethanol. After each step, excess liquid was wicked away with filter paper. Grids were coated with 100µg/ml poly-D-lysine (PDL) for 1h at room temperature. The excess of the liquid was wicked off. EVs were added to the grid and incubated overnight at 4°C in a moist chamber. The excess sample was wicked away using filter paper, and grids were subsequently stained with 2% phosphotungstic acid, pH 6.1 for 30 sec and allowed to air-dry overnight. Grids were imaged on a Tecnai T12 TEM (100kV LaB6 source), using an AMT CR41-S side-mounted 2K x 2K CCD camera..

### Purification of cellular fibronectin

Parental hMFs were cultured in complete medium overnight, and the medium was replaced with Opti-MEM after washing with PBS, 2 times. After 48 h, the conditioned medium was collected by centrifugation at 300 x *g* for 10 min. The supernatant was centrifuged at 100,000 x *g* for 90 min to pellet protein aggregates and EVs, including exosomes. The EV-depleted supernatant was incubated with Gelatin Sepharose 4B beads (GE Healthcare, 17-0956-01) at 4°C overnight on a rotator. The beads were washed with PBS, and cellular fibronectin was eluted with 8 M Urea. The eluted fractions were pooled and serially dialyzed in 4 M Urea in PBS, 1 M Urea in PBS, and CAPS/saline buffer (10 mM CAPS, pH 11.0, 1 mM CaCl_2_, 150 mM NaCl), 3 times.

### Preparation of colloidal gold cellular fibronectin

Colloidal gold beads of 16 nm (pH 7.0) (generous gift from Dr. Evan Krystofiak) were added to 10 µg of purified cellular fibronectin, mixed by inverting, and incubated at room temperature for 5 min. Unbound areas on the gold beads were blocked with 1% of PEG-8000 (final concentration of 0.05%) by mixing and centrifuging at 15,000 x *g* for 10 min at 4°C. The supernatant was removed, and the soft pellet obtained was resuspended in 0.1 M HEPES, pH 7.4.

### *In vitro* purified component assay for fibronectin assembly

1 μg of purified cellular fibronectin or plasma fibronectin (Sigma Aldrich F1056) was mixed with 1×10^8^ of purified EVs in Dulbecco’s PBS with calcium and magnesium (+/- 500 µM Mn^2+^ or +/- 5 mM EDTA) in pre-blocked tubes with 5% BSA in PBS and incubated for 72h in a PCR block at 37°C with a heated lid to prevent evaporation of the reaction. The reactions were plated on poly-D-lysine (100µg/ml) coated glass coverslips and incubated overnight at 4°C. The coverslips were washed briefly with PBS, fixed with 4% PFA for 15 min at room temperature, and blocked for 1h at room temperature with 5% BSA in PBS. The samples were immunostained for fibronectin and Tsg101, Hsp70, or CD63. For the staining of Tsg101 and Hsp70, the primary antibodies were diluted in blocking buffer supplemented with 0.1% Triton X-100.Images were acquired using a Nikon Eclipse TE2000E wide-field microscope equipped with a Nikon Plan Apo 100X/1.40 oil immersion objective lens and a CCD camera described above or using a Nikon A1R-HD25 confocal microscope equipped with a Nikon Apo TRIF 60x/1.49 oil immersion objective lens.

Images were thresholded and quantified using ImageJ. For integrin blocking assay, normal mouse IgG (Santa Cruz Biotechnology sc-2025), P1H4, an integrin α4 blocking antibody (Millipore MAB16983Z), P1D6, an integrin α5 blocking antibody (Santa Cruz Biotechnology sc- 13547), LM609, an integrin αvβ3 blocking antibody (Millipore MAB1976), and P1F6, an integrin αvβ5 blocking antibody (Santa Cruz Biotechnology sc-81632 L) were purchased. For the DOC) solubility assay, the reactions incubated for 72h were added to the same volume of 2X DOC lysis buffer (4% sodium DOC, 40 mM Tris-HCl, pH 8.8, 4 mM phenylmethylsulfonyl fluoride (PMSF), 4 mM ethylenediaminetetraacetic acid (EDTA), 4 mM iodoacetic acid, 4 mM N-ethylmaleimide (NEM)) and incubated for 5 min at room temperature. Samples were spun at 15,000 x *g* for 30 min at 22°C. The supernatant was collected as a DOC-soluble fraction, and the pellet was resolved with 2X Laemmli sample buffer as a DOC-insoluble fraction. Both DOC-soluble and -insoluble fractions were applied to Western blot probed for fibronectin.

### Scanning Electron Microscopy

*In vitro* assembled fibronectin was carefully plated on 100 µg/ml PDL-coated glass coverslips and incubated at 4°C overnight. The samples were fixed in 2.5% glutaraldehyde in 0.1 M cacodylate buffer at room temperature for 1h, washed with 0.1M cacodylate buffer (no Ca^++^). Samples were sequentially postfixed in 1% tannic acid in 0.1 M cacodylate buffer at room temperature for 20 min, and 0.05% osmium tetroxide (OsO_4_) for 30 min and with 1% uranyl acetate for 30 min. Subsequently, the samples were dehydrated through a graded ethanol series (25%,50%,75%, 80%,85%, 90%, and 95%) for 5 min each and in 100% ethanol for 15 min, 3 times. The coverslips were dried in a critical point dryer (Tousimis Samdri-PVT-3D) and mounted on an aluminum stub with carbon adhesive tab and sputter coated with gold palladium using the Cressington Sputter Coater 108. The samples were imaged using the FEI Q-250 scanning electron microscope at 10 kV of accelerating voltage.

### Co-immunoprecipitation of protein complexes in small EVs

The protocol adapted and modified from Zimmerman et al. ^59^ was used to co-precipitate syndecan-1 and its protein complexes in small EVs. Briefly, small EVs isolated from hMF overexpressing mouse syndecan-1-mScarlet were extracted with lysis buffer (50 mM 2-(*N*-morpholino) ethanesulfonic acid (MES), pH 6.8, 70 mM NaCl, 1 mM EGTA, 1 mM MgCl2, 1% NP-40, EDTA-free protease inhibitors cocktail) and cleared by centrifugation at 12,000 rpm for 10 min. The cleared extract was mixed with the ChromoTek RFP-Trap® Magnetic Agarose (Proteinteck rtma) and incubated overnight at 4°C by rotation. The beads were resuspended with heparitinase buffer (50 mM HEPES, pH 6.5, 150 mM NaCl, 50 mM NaOAc, 5 mM CaCl2, 2 U/ml Heparinase I and III Blend (Sigma Aldrich H3917), 0.1 unit/ml Chondroitinase ABC (Sigma Aldrich C3667)) and incubated at 37°C. After 4h of incubation, the protein complexes were released from the beads by boiling for 5 min in 2X Laemmli sample buffer and further used for Western blot.

### Mass spectrometry

Large and small EVs isolated from hRFs were solubilized by an equal volume of 2X lysis buffer (20 mM HEPES, 300 mM NaCl, 4% NP-40, and 1% Sodium DOC). Lysed EVs were sonicated with Diagenode Bioruptor Standard Waterbath Sonicator UCD-200 in ice-cold water for 15 min (30 sec on and off). Cleared lysates were collected after centrifugation at 15,000 rpm for 30 min at 4°C. The protein concentration was determined using Pierce Micro BCA Assay (Thermo Fisher Scientific 23235). Protein samples were precipitated, reduced, alkylated, and digested with trypsin as described in Jimenez *et al*^60^. Quantitative proteomics analysis was performed using TMT Isobaric Mass Tagging reagents (Thermo Fisher Scientific 90066) by Vanderbilt University Proteomics Core. According to the manufacturer’s instructions, peptides were labeled with TMT reagents with each reconstituted protein sample being labeled with an individual vial of 0.8 mg TMTsixplex™ reagent. After labeling was complete and quenched, labeled peptides from each sample were combined. Fractionation was performed on the combined mixture using the Pierce High pH Reversed-Phase Peptide Fractionation Kit (Thermo Fisher Scientific 84868), using the manufacturer’s recommended protocol for TMT-labeled peptides. Elution steps consisted of the following: 10%, 12.5%, 15%, 17.5%, 20%, 22.5%, 25%, and 50% acetonitrile with 0.1% triethylamine. Eluted fractions were dried in a SpeedVac concentrator, and peptides were reconstituted in 0.2% formic acid for LC-MS/MS analysis using methods similar to those applied to tagged-SDC1. Peptides were gradient-eluted at a flow rate of 400 nL/min, using a 140-minute gradient. The gradient consisted of the following: 5-30 %B in 115 min, 30-50 %B in 10 min, 50-70 %B in 5 min, 70 %B for 2 min; 70-2%B in 1 min, followed by column equilibration at 2 %B. For the final peptide fraction, peptides were eluted from the reverse phase column using a gradient of 5-50 %B in 110 min, followed by 50-95 %B in 15 min, 95 %B for 4 min, 95-2 %B in 1 min, and 2 %B for 10 min. Peptides were analyzed on a Q Exactive Plus mass spectrometer (Thermo Fisher Scientific), equipped with a nanoelectrospray ionization source, using a data-dependent top15 method. Acquisition method parameters included MS1 with an MS AGC target value of 3e6, followed by 15 MS/MS scans of the most abundant ions detected in the preceding MS scan. The MS2 AGC target was set to 1e5. Dynamic exclusion was 20s, HCD collision energy was set to 29 nce, and peptide match and isotope exclusion were enabled.

For identification of peptides, HCD tandem mass spectra were searched in Proteome Discoverer 2.1 (Thermo Fisher Scientific) using SequestHT for database searching against a subset of the UniprotKB protein database containing *Homo sapiens* protein sequences. Parameters were based on factory default templates in Proteome Discoverer for processing and consensus workflows, which included the PWF_QE_Reporter_Based_Quan_SequestHT_Percolator processing workflow and the CWF_Comprehensive_Enhanced Annotation_Quan_Results export consensus workflow. Search parameters included trypsin cleavage rules with two missed cleavage sites, carbamidomethyl (C) and TMTsixplex (K, N-terminus) as static modifications, and a dynamic modification of oxidation (M). Percolator validation was performed with a target false discovery rate (FDR) setting of 0.01 for high confidence protein identifications. In the consensus workflow, an average reporter S/N threshold of 10 was applied, and no scaling or normalization modes were selected. Log2 protein ratios calculated in Proteome Discoverer were then fit to a normal distribution using non-linear (least squares) regression. The mean of the normal curve was used to normalize log2 protein ratios, and the mean and standard deviation of the normalized log2 ratios were used to calculate *p* values as previously described^61^. Subsequently, *p* values were corrected for multiple comparisons by the Benjamini-Hochberg (B-H) method^62^. Statistically significant changes were determined using the B-H method with an alpha (α) level 0.05 (<5% FDR)^62^.

To identify protein complexes immunoprecipitated with mouse syndecan-1 (described above), the sample was brought to a final concentration of 5% SDS, reduced with 10 mM DTT, and alkylated with 20 mM iodoacetamide. Proteins were then prepared for digestion on a S-Trap^TM^ micro spin column (ProtiFi) following manufacturer’s instructions. Aqueous phosphoric acid was added to a final concentration of 2.5%, followed by the addition of 90% methanol containing 100 mM TEAB (S-Trap binding/wash buffer) at 6X the volume. The sample was loaded onto the S-Trap column and centrifuged at 4,000 x *g*. The column was washed 4 times with binding/wash buffer. Proteins were digested with trypsin (Promega) in 50 mM TEAB, pH 8.0, for 1h at 47°C. Peptides were recovered by sequential elution with 40 µL each of 50 mM TEAB, 0.2% formic acid, and 0.2% formic acid in 50% acetonitrile. Eluted peptides were dried in a speed-vac concentrator, reconstituted in aqueous 0.2% formic acid, and analyzed by LC-coupled tandem mass spectrometry (LC-MS/MS) as described in Howard et al.^63^. Peptides were loaded onto a reverse phase capillary column using a Dionex Ultimate 3000 nanoLC and autosampler. The mobile phase solvents consisted of 0.1% formic acid, 99.9% water (solvent A) and 0.1% formic acid, 99.9% acetonitrile (solvent B). Peptides were gradient-eluted at a flow rate of 350 nL/min, using a 60-min gradient. The gradient consisted of the following: 0-46 min, 2-35% B; 46-50 min, 35-90% B; 50-51 min, 90% B; 51-52 min, 90-2% B; 52-60 min, 2% B. Peptides were analyzed using a data-dependent method on an Orbitrap Exploris 480 mass spectrometer (Thermo Scientific). The instrument acquisition method consisted of MS1 (AGC target value of 3×10^6^), followed by up to 15 MS/MS scans (AGC target of 1×10^5^) of the most abundant ions per MS scan. The intensity threshold for MS/MS was set to 1×10^4^, HCD collision energy was 30 nce, and dynamic exclusion (20sec) was enabled. Tandem mass spectra were searched with Sequest (Thermo Fisher Scientific) against a *Homo sapiens* database and mouse syndecan-1 created from the UniprotKB protein database appended with the mScarlet-tagged mouse syndecan-1 protein sequence . Variable modifications included +15.9949 on Met (oxidation) and +57.0214 on Cys (carbamidomethylation). Search results were assembled in Scaffold 5.1.0 (Proteome Software) using a minimum filtering criteria of 95% peptide probability and 99% protein probability.

### Super-resolution direct stochastic optical reconstruction microscopy (dSTORM) using the Oxford Nanoimager (ONi)

Purified small EVs were captured on a 100 µg/ml PDL-coated assay chip provided in the EV Profiler 2 kit (ONi) for 75 min, fixed with Fixative for 10 min, permeabilized with 0.01% Triton X-100 (Amresco M236-10ML-5pk) for 10 min, and blocked with Staining Buffer provided in the EV Profiler 2 kit for 1 h. Small EVs were incubated with CF^TM^ 568-labeled antibody against integrin α4 (P1H4), α5 (P1D6), αvβ5 (P1F6), activated integrin β1 (12G10, Biotium BNC682905-100), or activated integrin β3 (AP-5, kerafast EBW107), or Alexa Fluor 555-conjugated integrin αvβ3 antibody (Millipore MAB1976-AF555) with CF^TM^ 647-labeled SDC1CT antibody (Abnova PAB9567) and CF^TM^ 488-labeled CD63 antibody (ONi 800-00056) for 1 h and postfixed with Fixative for 10 min. The fluorophore-labeled antibodies which don’t show ordering information in this section were conjugated with CF^TM^ 568 succinimidyl ester (Sigma Aldrich SCJ4600027-1UMOL) or CF^TM^ 647 succinimidyl ester (Sigma Aldrich SCJ4600048-1UMOL) using the manufacturer’s instructions. For imaging, BCubed imaging buffer was applied to the chip. Images were acquired using a 100x oil objective lens equipped on the ONi Nanoimager with 3 channels (640 nm, 561 nm, and 488 nm lasers) and analyzed using a web-based software, CODI (alto.codi.bio). Protein count per EV was measured by dividing the number of localizations of individual fluorescent probes on single EVs by the median value of localizations for a single fluorescent reporter using the single extracellular vesicle Nanoscopy (SEVEN) assay described by Saftics *et al*^64^.

### Numbers and Statistics

The figure legends list the *n* values and independent experiment numbers for both quantitated data and representative images from experiments. All datasets were tested for normality using GraphPad Prism’s Kolmogorov-Smirnov or Shapiro-Wilk normality test. The two-sided unpaired Mann-Whitney or two-sided paired Wilcoxon test compared nonparametric data groups. Parametric data were compared using a two-sided unpaired or paired Student’s *t*-test. Scatter plots were plotted with median and interquartile range or mean ± standard error of the mean. Bar graphs were plotted as mean ± standard error of the mean. All graphs were created using Excel or GraphPad Prism.

### Use of AI

ChatGPT, version 4o, was used to draft parts of the Discussion, followed by editing.

## Acknowledgements

Funding was provided by NIH grants R01CA206458 (AMW and SMP), R01CA249684 (AMW), R01CA249424 (AMW), R50CA283661 (BHS), and AHA fellowship 19POST34370067 (ME). Electron microscopy was done in part through the Vanderbilt Cell Imaging Shared Resource, supported by NIH grants CA68485, DK20593, DK58404, DK59637, and EY08126.

---

**Supplementary Data 1:** Raw and analyzed TMT proteomics data, comparing SEVs to LEVs.

**Supplementary Data 2:** Proteomics analysis of SEV proteins coprecipitated with Syndecan-1-mScarlet.

**Extended Data Figure 1.**
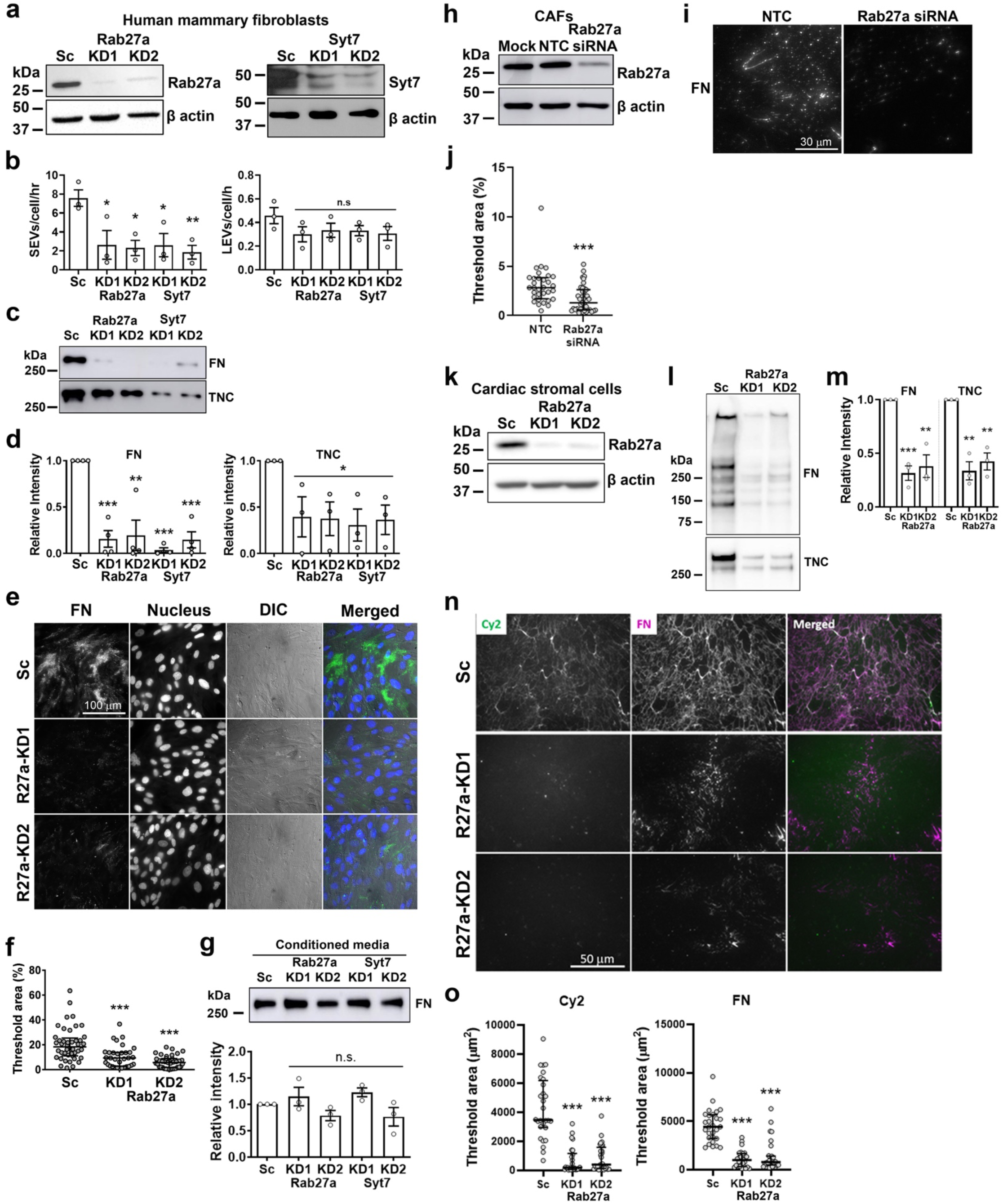
Exosome secretion is required for stromal matrix deposition. a, Western blots showing the knockdown expression of Rab27a and synaptotagmin VII (Syt7) in hTERT-immortalized human mammary fibroblasts (hMFs). Sc, scrambled control; KD, knockdown. **b,** EV secretion rates measured by nanoparticle tracking analysis (mean ± SEM). SEV, small extracellular vesicle; LEV, large extracellular vesicle. n = 3. *, *p* < 0.05; **, *p* < 0.01; n.s., not significant compared to Sc by student *t* test. **c,** Western blots showing ECM deposition in the cell-derived stromal matrix. FN, fibronectin; TNC, tenascin C. **d,** Quantitation of **c** relatively compared the band intensity to Sc (mean ± SEM). n = 4 for FN blots and n = 3 for TNC blots. **e,** Representative total internal reflection fluorescence micrographs of hMFs immunostained for FN. DIC, differential interference contrast. **f,** Integrated intensity of **e** (median with interquartile range). n ≥ 31 images from 3 independent experiments. **g,** Western blots for fibronectin in conditioned media collected from hMFs cultured in serum-free Opti-MEM and relative intensity of the FN bands compared to Sc (mean ± SEM). n = 3. **h,** Western blots showing transient knockdown of Rab27a in cancer-associated fibroblasts (CAFs). NTC, non-targeting control. **i,** CAF-derived stromal matrix immunostained for FN. **j,** Integrated intensity of **i** (median with interquartile range). n ≥ 32 images from 2 independent experiments. **k,** Western blots showing the knockdown expression of Rab27a in mouse cardiac stromal cells (CSCs). **l,** Western blots for FN and TNC from CSC-derived stromal matrix. **m,** Relative intensity of FN and TNC compared to scrambled controls (mean ± SEM) from **l**. n = 3. **n,** Immunostaining images of CSC-derived stromal matrix for the whole matrix (Cy2) and FN. **o,** Integrated intensity of **n** (median with interquartile range). n ≥ 26 from 3 independent experiments. n.s., not significant; *, *p* < 0.05; **, *p* < 0.01; ***, *p* < 0.001 compared to Sc determined by two-sided Student’s *t*-test for bar graphs or Mann-Whitney *U* test for scatter plots.

**Extended Data Figure 2.**
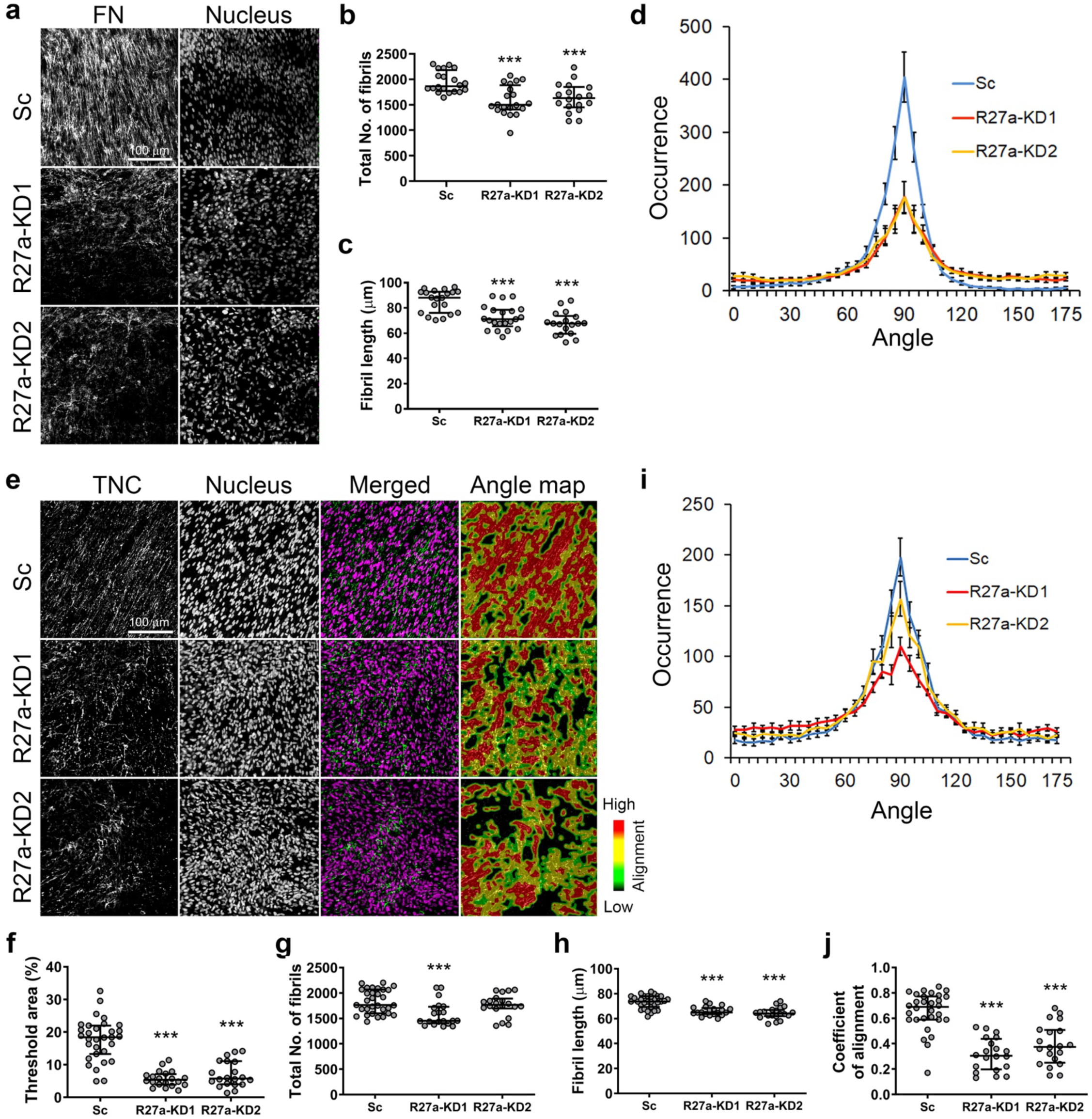
Exosome secretion is required for matrix assembly and alignment. a, Representative confocal images by maximum intensity projection of fibronectin (FN) immunostaining and nucleus staining related to Fig 1f. Scrambled control (Sc), Rab27a-knockdown (R27a-KD). **b – d,** Total number (**b**), length (**c**), and occurrence of aligned fibers with respect to 90° angle (**d**) of FN fibrils. n ≥ 18 images from 4 independent experiments (median with interquartile range for **b** and **c**, mean ± SEM for **d**). **e,** Representative confocal images by maximum intensity projection of tenascin C (TNC) immunostaining from Sc or Rab27a-KD hMFs cultured in collagen gels for 7 days. **f – j,** Threshold area (**f**), total number (**g**), length (**h**), occurrence of aligned fibers with respect to 90° angle (**i**), and coefficient of alignment (**j**) of TNC fibrils from **e**. n ≥ 20 images from 4 independent experiments (median with interquartile range for **f – h** and **j**, mean ± SEM for **i**). ***, *p* < 0.001 compared to Sc determined by Mann-Whitney *U* test.

**Extended Data Figure 3.**
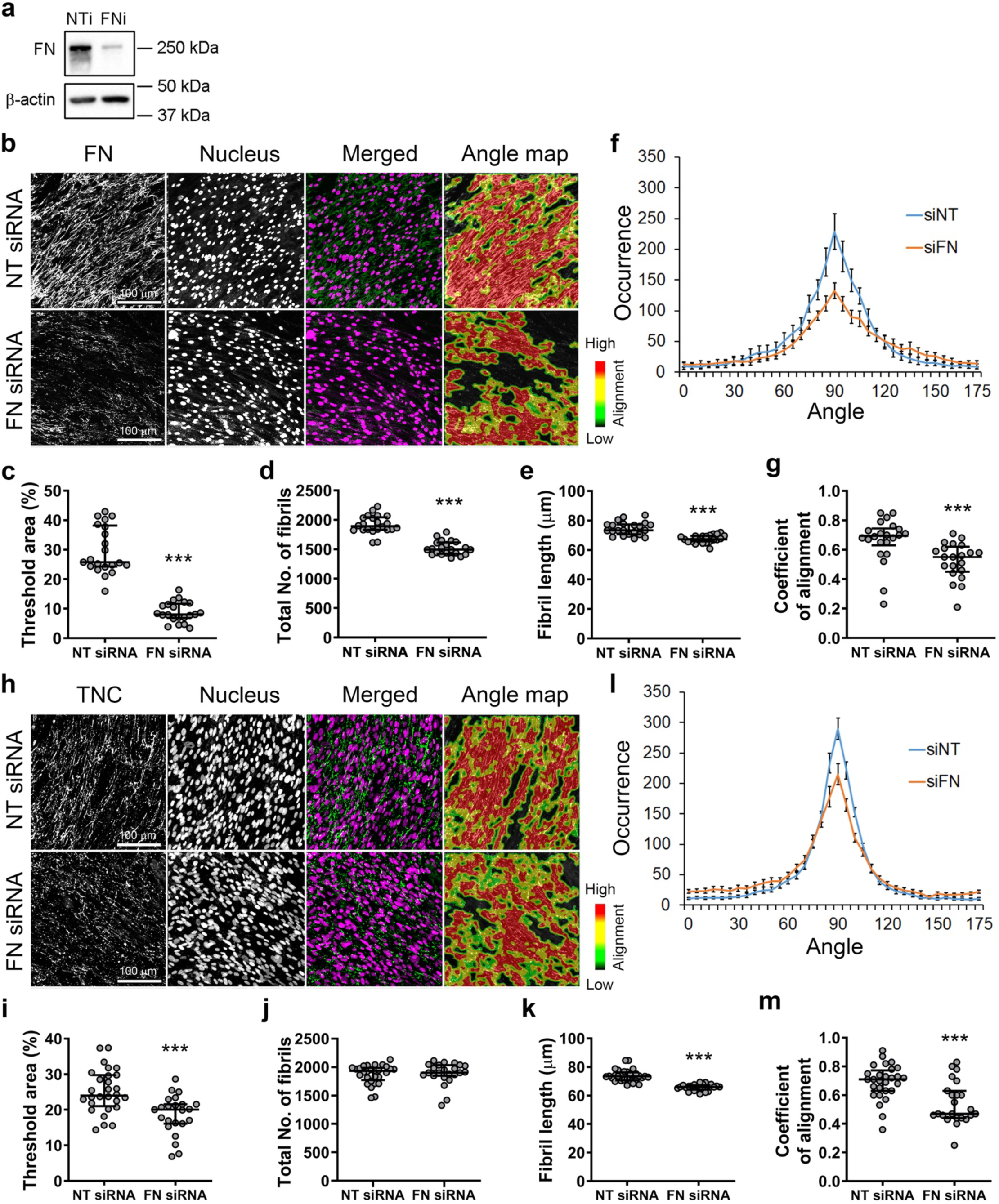
Fibronectin expression is critical for matrix assembly and alignment. a, Western blots for the transient knockdown of fibronectin (FN) in hMFs. NTi, non-targeting siRNA. FNi, FN siRNA. **b,** Representative confocal images by maximum intensity projection of FN immunostaining from NTi or FNi hMFs cultured in collagen gels for 7 days. **c – g,** Threshold area (**c**), total number (**d**), length (**e**), occurrence of aligned fibers with respect to 90° angle (**f**), and coefficient of alignment (**g**) of FN fibrils from **b**. n = 21 images from 3 independent experiments (median with interquartile range for **c – e** and **g**, mean ± SEM for **f**). **h,** Representative confocal images by maximum intensity projection of tenascin C (TNC) immunostaining from NTi or FNi hMFs cultured in collagen gels for 7 days. **i – m,** Threshold area (**i**), total number (**j**), length (**k**), occurrence of aligned fibers with respect to 90° angle (**l**), and coefficient of alignment (**m**) of TNC fibrils from **h**. n ≥ 23 images from 3 independent experiments (median with interquartile range for **i – k** and **m**, mean ± SEM for **l**). ***, *p* < 0.001 compared to NT siRNA determined by Mann-Whitney *U* test.

**Extended Data Figure 4.**
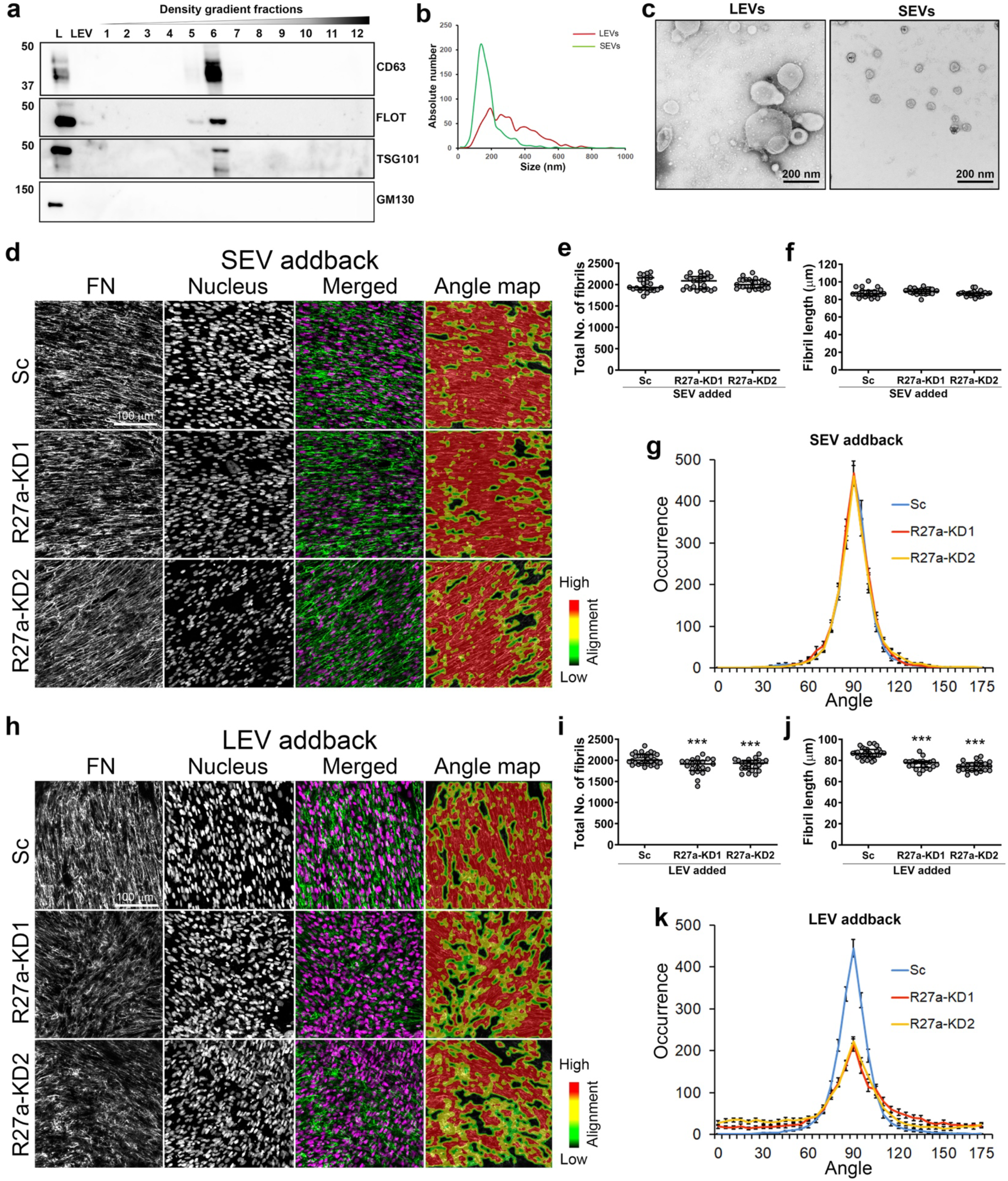
Purified SEVs/exosomes can rescue matrix alignment. a, Western blots to verify EV purification. L, cell lysate. LEV, large EVs. FLOT, flotillin. **b,** Nanoparticle tracking analysis of purified EVs. LEVs, large EVs. SEVs, small EVs. **c,** Representative negative staining transmission electron micrographs of purified LEVs and SEVs. **d,** Representative confocal images by maximum intensity projection of FN immunostaining from Sc or Rab27a-KD hMFs cultured in collagen gels with purified large EVs (LEVs) for 7 days related to Fig 1f. **e – g,** Total number (**e**), length (**f**), and occurrence of aligned fibers with respect to 90° angle (**g**) of FN fibrils from **d** (median with interquartile range for **e** and **f**, mean ± SEM for **g**). n ≥ 21 images from 3 independent experiments. **h,** Representative confocal images by maximum intensity projection of FN immunostaining from Sc or Rab27a-KD hMFs cultured in collagen gels with purified small EVs (SEVs) for 7 days related to Fig 1f. **i – k,** Total number (**i**), length (**j**), and occurrence of aligned fibers with respect to 90° angle (**k**) of FN fibrils from **h** (median with interquartile range for **i** and **j**, mean ± SEM for **k**). n ≥ 22 images from 3 independent experiments. ***, *p* < 0.001 compared to Sc determined by Mann-Whitney *U* test.

**Extended Data Figure 5.**
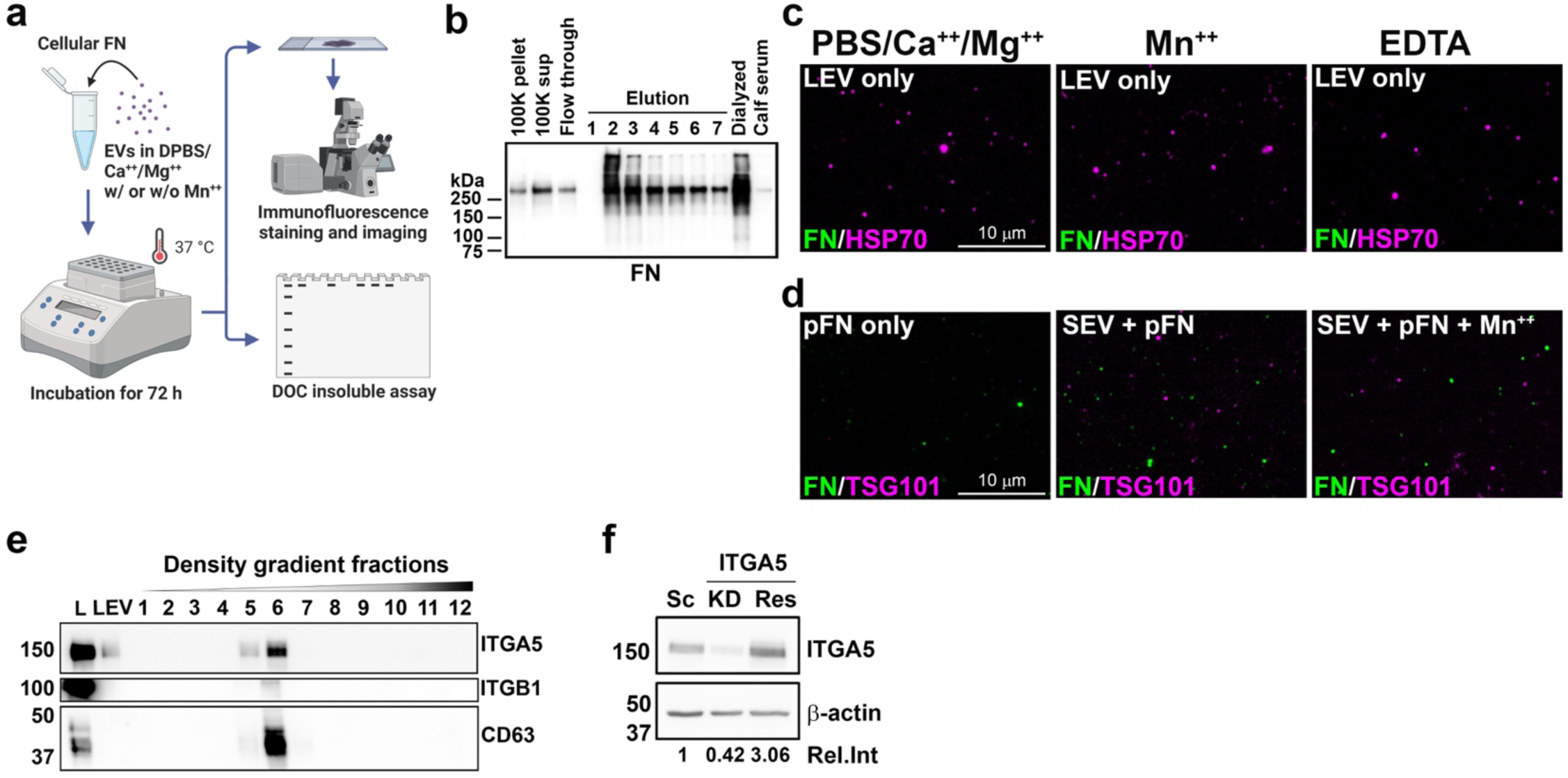
Integrin. α**5 promotes fibronectin assembly. a,** A schematic diagram showing the procedure of cell-free purified component assay for fibronectin assembly. Created with BioRender.com. **b,** Western blots of cellular fibronectin purification from the conditioned media of hMFs. 100K, 100,000 x *g* ultracentrifugation. **c,** Representative wide-field images for cell-free purified component assay for fibronectin assembly with large EV (LEV) only immunostained with FN and HSP70 for LEVs in the presence of 500 μM MnCl_2_ or 5 mM EDTA in PBS/Ca^++^/Mg^++^ related to Fig 3a. **d,** Representative wide-field images for cell-free purified component assay for fibronectin assembly with plasma fibronectin (pFN) by parental small EVs (SEVs) immunostained with FN and TSG101 for SEVs in the presence of 500 μM MnCl_2_ in PBS. **e,** Western blots for integrins α5 (ITGA5) and β1 (ITGB1) in EVs purified from the conditioned media of hMFs. L, cell lysate. **f,** Western blots for the manipulation of integrins α5 expression in hMFs. Sc, scrambled. KD, knockdown. Res, rescued. Rel. Int., relative intensity.

**Extended Figure 6.**
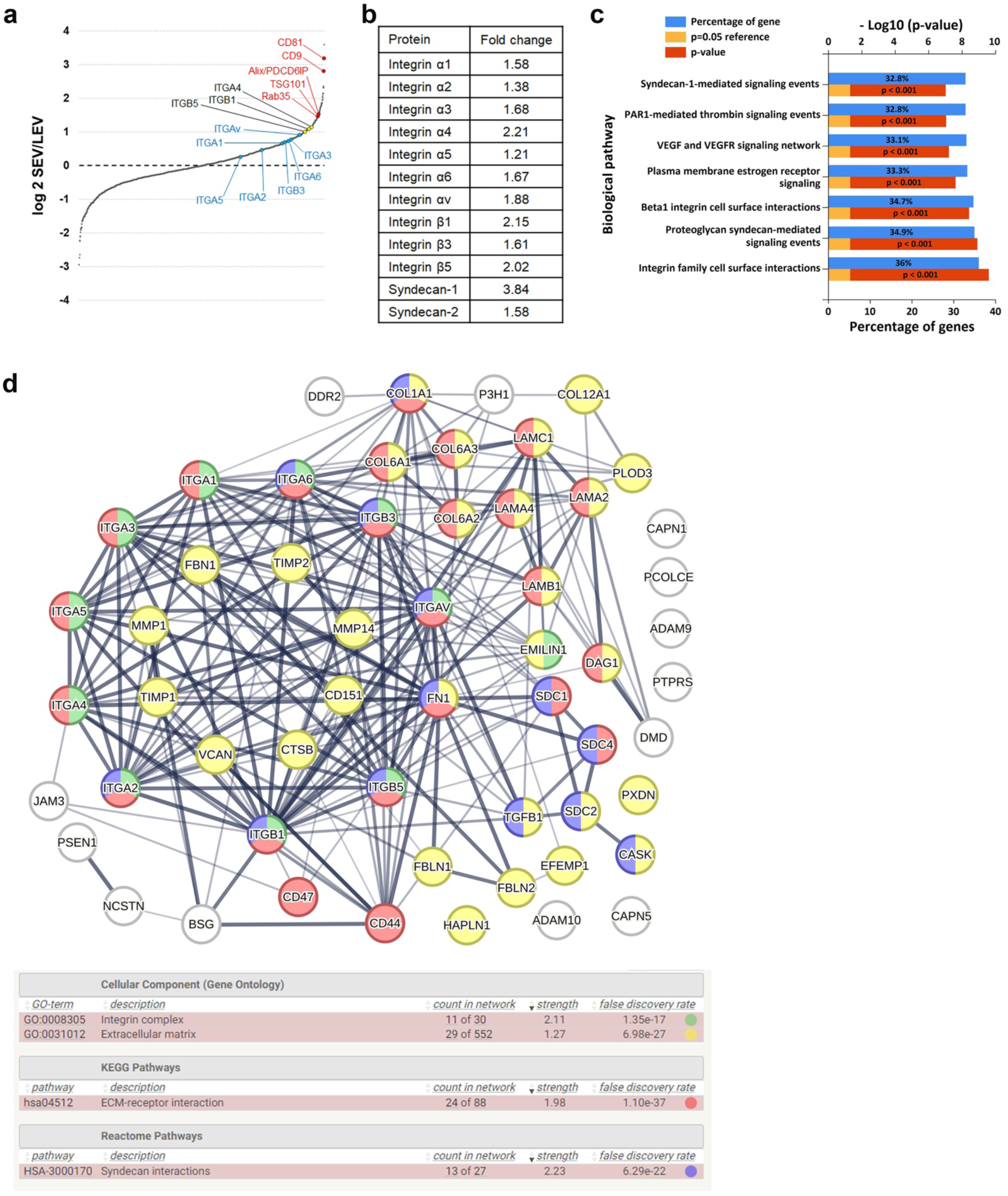
Integrins and syndecans are abundant in small EVs. a, Protein rank plot showing the abundance of proteins by the magnitude of change between small EVs (SEV) and large EVs (LEV). Red color indicates tetraspanins and SEV biogenesis molecules >2-fold enriched in SEVs compared to LEVs. Black color indicates integrins >2-fold enriched in SEVs compared to LEVs. Blue color indicates integrins enriched but not >2-fold in SEVs compared to LEVs. **b,** Fold change of the gene expression level of integrins and syndecans in SEVs compared to LEVs. **c,** Biological pathway of the top 7 gene enrichment from proteomics analysis. **d,** STRING protein-protein interaction network of extracellular matrix-related proteins abundant in small EVs.

**Extended Figure 7.**
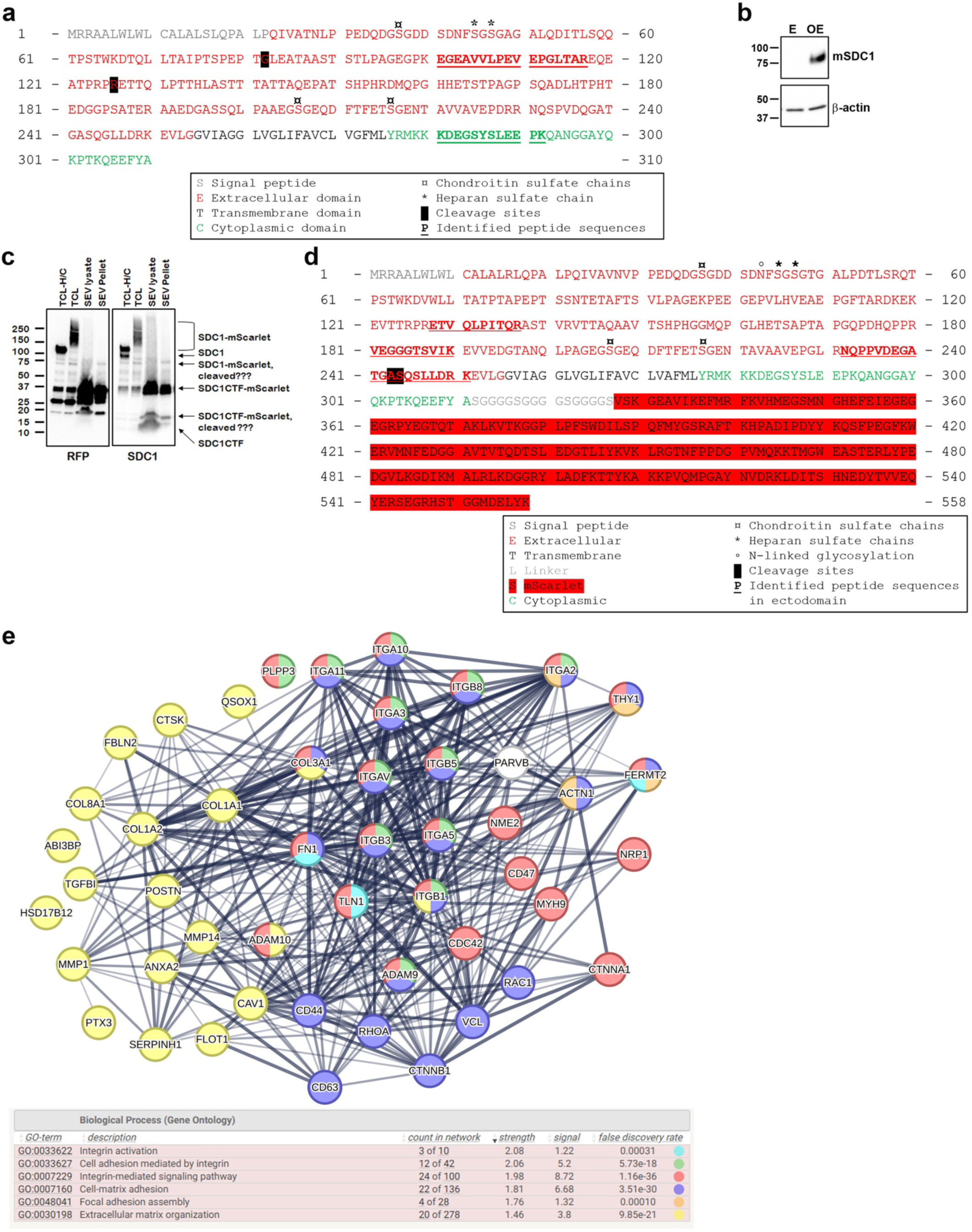
Syndecan-1 is presented in small EVs as multiple fragments. a, Amino acid sequence of human syndecan-1. Domains and identified peptides from hMF SEV proteomics are indicated in the boxed legend. **b,** Western blots of mouse syndecan-1 (mSDC1) in empty vector-overexpressed (E) and mSDC1-overexpressed (OE) fibroblasts. **c,** Western blots of immunoprecipitation experiment for mouse syndecan-1-mScarlet using RFP-Trap^®^, probed for RFP or SDC1. TCL, total cell lysate. H/C, heparinase-chondroitinase-treated. SEV, small EV. SDC1, syndecan 1. CTF, cytoplasmic tail fragment. **d,** Amino acid sequence of mouse syndecan-1 tagged with mScarlet. Domains and identified peptides from the proteomics of mouse syndecan-1-mScarlet pulldown from SEVs are indicated in the boxed legend. **e,** STRING protein-protein interaction network of extracellular matrix-related proteins present in the mouse syndecan-1-mScarlet pulldown from SEVs.

**Extended Figure 8.**
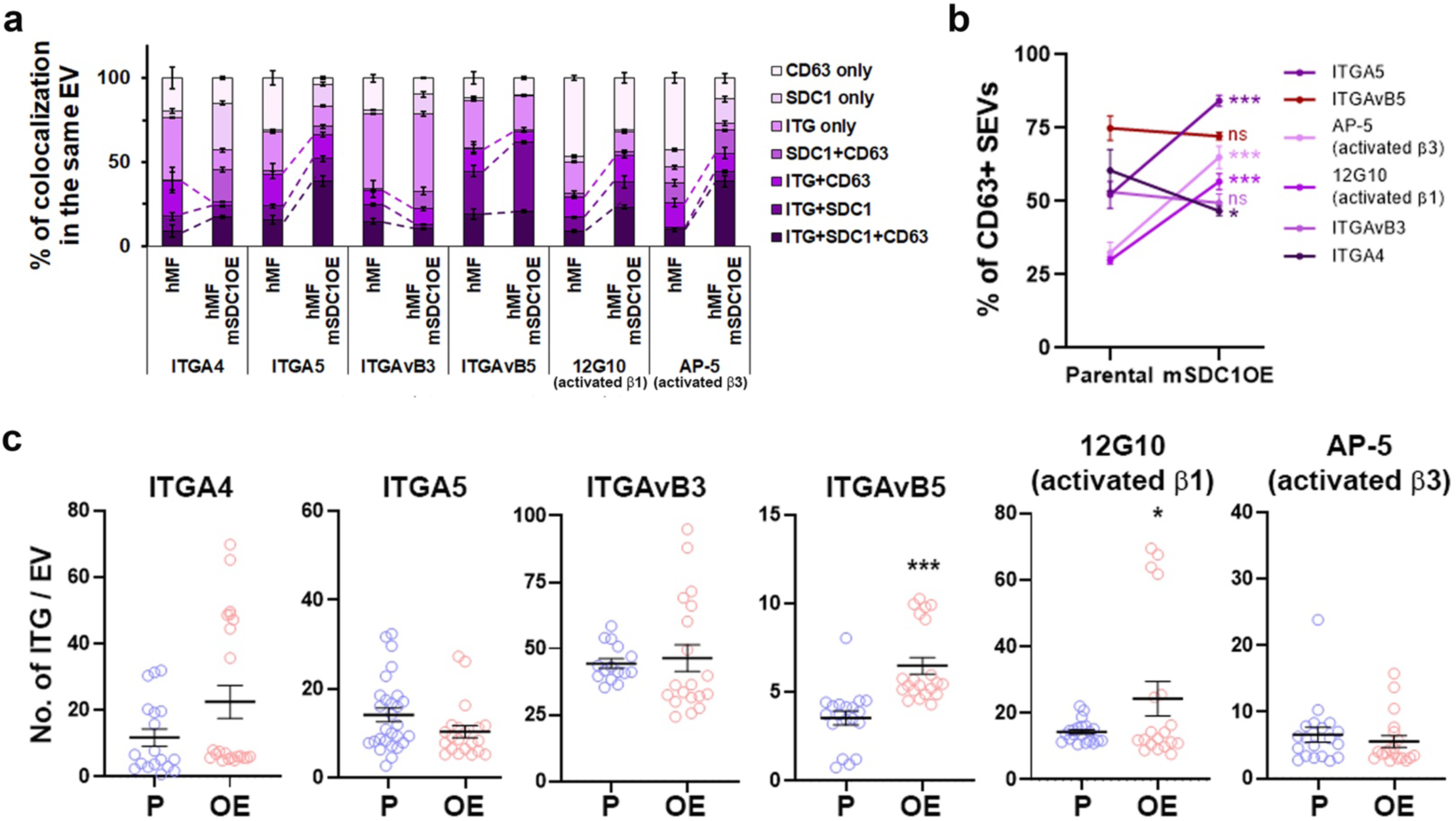
Integrins and syndecan-1 are colocalized in the same small EVs. a, Percentage of colocalization of integrins, syndecan-1 (SDC1), and CD63 in the same SEVs quantified from dSTORM images (mean ± SEM). Dot lines indicate the difference in percentage of integrins colocalized with SDC1, CD63, or both in the same small EVs purified from parental and mouse SDC1-overexpressing hMFs. **b,** Percentage of integrins in small EVs containing CD63 (mean ± SEM). n ≥ 15 field-of-views from 3 or 4 independent experiments. n.s., not significant; *, *p* < 0.05; ***, *p* < 0.001 compared to parental small EVs determined by unpaired *t*-test. **c,** Comparison of the number of integrins per EV in the population of integrin+syndecan-double positive small EVs purified from parental (P) and mouse SDC1-overexpressing (OE) hMFs. Mean ± SEM. Each dot represents a field-of-view (≥ 15) from 3 or 4 independent experiments. *, *p* < 0.05; ***, *p* < 0.001 compared to parental small EVs determined by unpaired *t*-test.

